# Structure-Based Design of a Highly Immunogenic, Conformationally Stabilized FimH Antigen for a Urinary Tract Infection Vaccine

**DOI:** 10.1101/2024.06.10.598184

**Authors:** Natalie C. Silmon de Monerri, Ye Che, Joshua A. Lees, Jayasankar Jasti, Huixian Wu, Matthew C. Griffor, Srinivas Kodali, Julio Caesar Hawkins, Jacqueline Lypowy, Christopher Ponce, Kieran Curley, Alexandre Esadze, Juan Carcamo, David Keeney, Arthur Illenberger, Yury V. Matsuka, Suman Shanker, Laurent Chorro, Alexey V. Gribenko, Seungil Han, Annaliesa S. Anderson, Robert G. K. Donald

## Abstract

Adhesion of *E. coli* to the urinary tract epithelium is a critical step in establishing urinary tract infections. FimH is an adhesin positioned on the fimbrial tip which binds to mannosylated proteins on the urinary tract epithelium via its lectin domain (FimH_LD_). FimH is of interest as a target of vaccines to prevent urinary tract infections (UTI). Previously, difficulties in obtaining purified recombinant FimH from *E. coli* along with the poor inherent immunogenicity of FimH have hindered the development of effective FimH vaccine candidates. To overcome these challenges, we have devised a novel production method using mammalian cells to produce high yields of homogeneous FimH protein with comparable biochemical and immunogenic properties to FimH produced in *E. coli.* Next, to optimize conformational stability and immunogenicity of FimH, we used a computational approach to design improved FimH mutants and evaluated their biophysical and biochemical properties, and murine immunogenicity. This approach identified a highly immunogenic FimH variant (FimH-DSG TM) that is produced at high yields in mammalian cells. By x-ray crystallography, we confirmed that the stabilized structure of the FimH_LD_ in FimH-DSG TM is similar to native FimH on the fimbrial tip. Characterization of monoclonal antibodies elicited by FimH-DSG TM that can block bacterial binding to mannosylated surfaces identified 4 non-overlapping binding sites whose epitopes were mapped via a combinatorial cryogenic electron microscopy approach. Novel inhibitory epitopes in the lectin binding FimH were identified, revealing diverse functional mechanisms of FimH-directed antibodies with relevance to FimH-targeted UTI vaccines.

**Author summary:** *Escherichia coli* is the primary cause of urinary tract infections. Adherence to uroepithelial surfaces is mediated by the pilus adhesin protein FimH, which is of interest as a vaccine candidate. We developed a method for producing recombinant FimH at bioprocess scale, previously a barrier to commercial development. Structure-based design and screening was used to identify a novel FimH vaccine candidate with improved stability and immunogenicity in mice. Structure of this full-length protein was determined by X-ray crystallography and shown to closely resemble the pilus adhesin present in its native form on the bacterial surface. Binding sites of biologically active FimH monoclonal antibodies were determined by X-ray crystallography or by cryo-electron microscopy, providing insights into mechanisms by which antibodies block binding of the bacteria to urinary tract receptors.

**One sentence summary:** Structure-based design of a conformationally stabilized *E. coli* FimH vaccine candidate capable of eliciting antibodies to diverse epitopes with the ability to block bacterial binding to bladder epithelial cells.

## Introduction

Uncomplicated urinary tract infections (UTI) affect approximately 50% of women at least once during their lifetime (1). In addition, sepsis-associated in-hospital mortality is significantly linked to UTI, resulting in a substantial burden on healthcare systems. Multidrug-resistant bacteria are frequently associated with UTI, which impacts routine urological practices (2). While several organisms can cause UTI, the most common agent is uropathogenic *Escherichia coli* (UPEC), which is associated with 75% of uncomplicated and 65% of complicated UTI (3). UPEC colonize the gastrointestinal tract and migrate from the fecal flora to the urogenital tract, where they adhere to host uroepithelial cells and establish a reservoir for ascending infections of the urinary tract (4). Adhesion is facilitated by fimbrial adhesins located on the bacterial surface, including type 1 fimbriae, which bind to mannosylated glycoproteins on bladder epithelial cells (5) as well as those secreted into the urine e.g. uromodulin (6).

Type 1 fimbriae are highly conserved among clinical UPEC isolates and are encoded by the *fim* gene cluster, which encodes chaperone and usher proteins (FimC, FimD), various structural subunits (FimA, FimF, FimG), and an adhesin called FimH that is displayed on the fimbrial tip (7). FimH is essential for all characteristics of UTI in mouse models that mimic aspects of human UPEC bladder infection: it mediates adhesion to target cells, intracellular invasion, biofilm formation, and resistance to killing by neutrophils (8–10). FimH is under positive selection in *E. coli* human cystitis isolates and positively selected residues have been proposed to influence virulence in mouse models of cystitis (11). In addition, it is important for uroepithelial cell shedding and invasion of bladder cells by *E. coli* (12, 13). Small molecule inhibitors that target FimH by mimicking mannosylated receptors are efficacious against UTI in animal models, further validating the role of FimH in UTI (14, 15).

FimH is composed of two domains, an N-terminal lectin binding domain (FimH_LD_) responsible for binding to the terminal mannose moiety on epithelial glycoproteins, and a C-terminal pilin domain (FimH_PD_). FimH_PD_ serves to link FimH to other structural subunits of the pilus such as FimG, through a mechanism called donor strand complementation (16). FimH_PD_ possesses an incomplete immunoglobulin-like fold, containing a groove that provides a binding site for the donor N-terminal β-strand of FimG, which when bound forms a strong intermolecular linkage with FimH. While FimH_LD_ can be expressed in a soluble, stable form, full-length FimH is unstable unless produced as a complex with the chaperone FimC or complemented with the FimG N-terminal donor strand peptide (as an exogenous peptide or directly fused to FimH via a polypeptide linker) (17, 18). FimH_LD_ transitions between two endpoint conformations: a ‘closed’ conformation with a high affinity for mannosylated proteins, and an ‘open’ conformation with a relatively compressed structure, a wide mannose binding pocket, that binds to mannosylated proteins with low affinity (19). Conformation and ligand-binding properties of FimH_LD_ are under the allosteric control of FimH_PD_, which itself exhibits only minor structural changes upon ligand binding (20, 21). Under static conditions, interaction between the lectin and pilin domains stabilizes FimH_LD_ in a low-affinity, open conformation (21, 22). Upon binding to a mannose derivative (mannoside) ligand, FimH_LD_ undergoes a conformational change leading to a high affinity state (with a 1000 to 100,000-fold higher affinity for mannose (21)), where the FimH_LD_ and FimH_PD_ remain in close contact. In the absence of negative allosteric regulation exerted by the FimH_PD_, isolated recombinant FimH_LD_ is locked in the high-affinity state and is highly stable (19). FimH is a key target for development of candidate vaccines to prevent UTI. The proposed primary mechanism of action of a FimH vaccine is inhibition of bacterial adhesion to urinary tract epithelial cells (23). Immunization with recombinant FimH_LD_ or full length FimH in complex with FimC (FimCH) is protective in both mouse and non-human primate models of UTI (23–32). Safety and immunogenicity of four doses of FimCH combined with a TLR4 adjuvant has been evaluated in humans in a Phase 1 study (33). In this exploratory analysis, female subjects with recurrent UTI had fewer UTI episodes following immunization, suggesting that the vaccine may reduce their frequency.

Development of FimH vaccines has been hindered significantly by two major issues: low yield of recombinant protein in *E. coli* and the requirement for multiple doses to elicit immune responses against *E. coli* (*33, 34*). In native *E. coli,* FimH is produced in the periplasm which facilitates disulfide bond formation (35). Typical yields of recombinant FimH reported at lab-bench scale are 3-5 mg/L for the purified FimCH complex and 4-10 mg/L for FimH_LD_ (18, 19); therefore, although the antigen has promise, it cannot be made in high enough quantities to proceed with clinical evaluation. In the current study, a mammalian expression platform was developed which yields high quantities of correctly folded FimH, providing a path forward to manufacture sufficient quantities of FimH to enable clinical trials.

To address the relatively poor immunogenicity of FimH, we used a structure-based design strategy to engineer FimH proteins predicted to exhibit superior immunogenicity compared to the wild type (WT) protein. It has been suggested that locking FimH_LD_ in the low affinity, ‘open’ conformation, induces the production of antibodies that can inhibit adhesion (36, 37). Conformational stabilization of antigens has been successful in improving immunogenicity and stability for various viral vaccine antigen candidates including Respiratory Syncytial Virus F protein (38–41). In this study, predicted conformationally stabilized FimH mutants were produced in mammalian cells and screened in a series of biochemical and biophysical assays followed by immunogenicity studies in mice. This screening approach led to the identification of a full length, donor strand-complemented FimH fusion antigen, FimH-DSG TM, which can be produced at a large scale in mammalian cells without a chaperone. By X-ray crystallography, we confirmed that the FimH_LD_ of this mutant was stabilized in an open conformation, similar to its native presentation on pili.

Conformers can have quite different structures wherein the accessibility (or conformation) of epitopes in one conformer versus another differs (42). Therefore, depending on which conformer is used for immunization, immune responses (quality, magnitude, specificity) can differ significantly, as observed in this study. For this reason, it was important to understand the nature of the inhibitory Mabs elicited by the FimH-DSG adhesin.

## Results

### Production of FimH in mammalian cells

The low yields of WT FimH_LD_ or FimCH produced in *E. coli* are well documented (18, 19). Given these challenges, we explored production of FimH using an Expi293 mammalian cell expression system (**Fig. 1A**), wherein the protein is secreted into the cell culture medium using a eukaryotic signal peptide. Fusion of FimH to the mouse IgGκ signal peptide yielded protein that was correctly processed at the N-terminus (confirmed by mass spectrometry (data not shown)). Non-native N-linked glycosylation commonly occurs during heterologous expression of proteins in mammalian cells and can interfere with biological function as well as antigenicity of recombinant proteins. Non-native N-linked glycosylation on FimH_LD_ at residues N7 and N70 was removed via mutation of target Asparagine residues to Serine, which was selected as a conservative substitution. WT and conformationally locked FimH_LD_ (V27C L34C, described previously (19)), harboring N7S and N70S mutations to remove glycosylation, were expressed in 20 ml Expi293 cultures, which yielded 17 mg and 10.8 mg protein respectively.

**Fig. 1.**
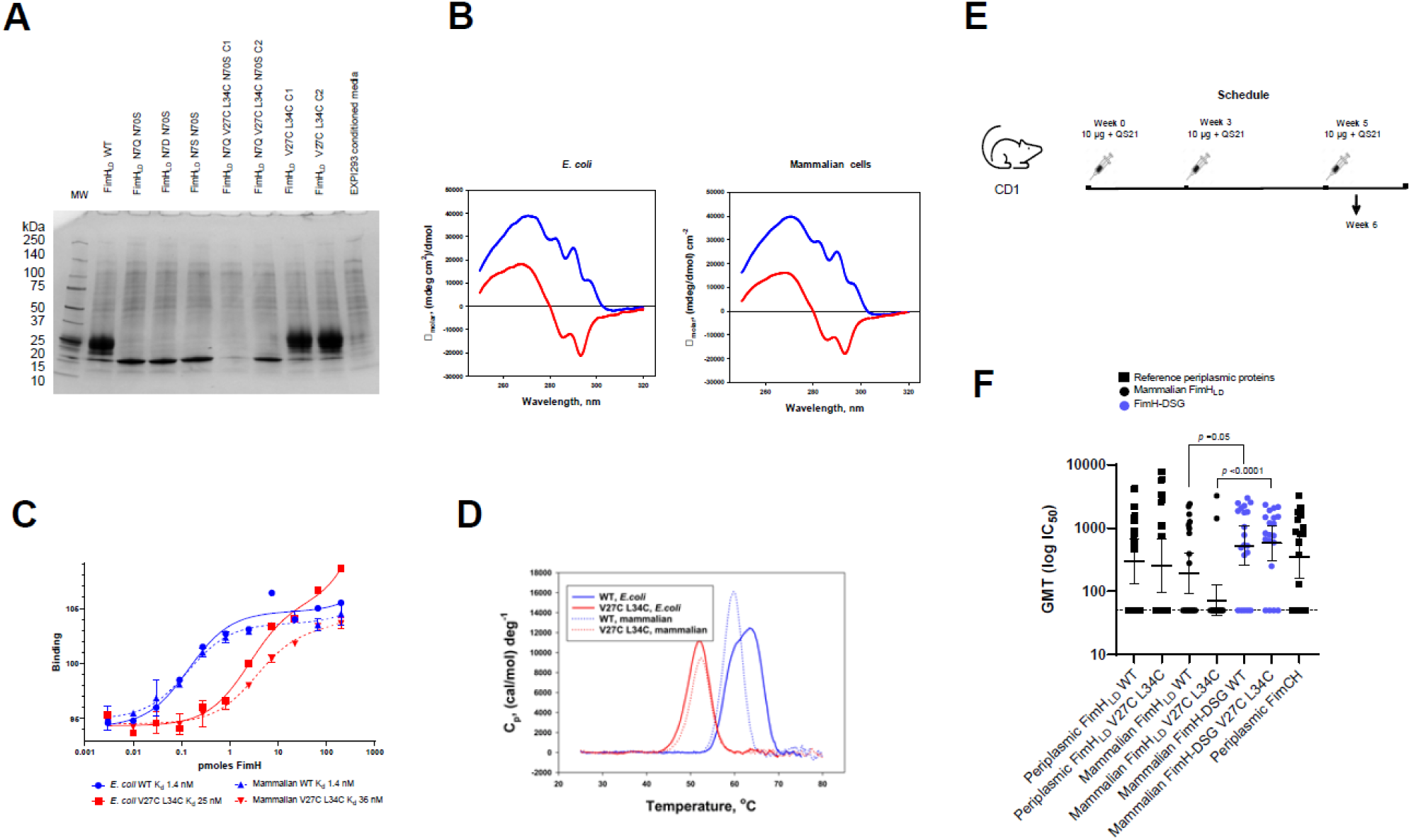
FimHLD produced in mammalian cells has comparable biophysical properties to material produced in *E. coli*. WT or conformation stabilized FimH_LD_ were expressed in Expi293 cells along with mutants designed to remove non-native glycosylation on residues N7 and N70. **(A)** Batch purified proteins run on SDS-PAGE gel stained with Coomassie blue. C1 and C2 designate two clones of FimH_LD_ N7Q V27C L34C N70S and FimH_LD_ V27C L34C C1 that were evaluated for expression. (**B**). Near-UV CD spectra of FimH_LD_ produced in *E. coli* and mammalian expression systems. Spectra of FimH_LD_ WT (blue) and V27C L34C (red) are shown. **(C)** Affinity of WT and V27C L34C FimH_LD_ produced in *E. coli* or mammalian cells for alpha-D-mannopyranoside by FP. **(D)** Thermal stability of WT and V27C L34C FimH_LD_ produced in *E. coli* or mammalian cells. **(E)** Mouse study design: CD-1 mice were immunized 3 times with 10 µg FimH proteins with QS21 adjuvant. **(F)** Sera were analyzed for the ability to block FimH-expressing *E. coli* binding to yeast mannan; bars represent geometric mean IC_50_ and 95% confidence intervals. Statistical significance (*p-*value) of differences in responses between groups was determined using an unpaired t-test with Welch’s correction applied to log-transformed data; the bars and asterisk illustrate the significance of the difference in response for comparisons.

Mammalian and *E. coli* derived FimH_LD_ WT and V27C L34C were further characterized using biophysical and biochemical assays. The *E. coli* and mammalian derived FimH_LD_ WT and mutant proteins had essentially identical near-UV circular dichroism (CD) spectra (**Fig. 1B**), indicating that tertiary structure of these proteins is very similar. Ligand binding affinity was determined using a direct binding fluorescence polarization (FP) assay with a fluorescein-conjugated octylbiphenylmannopyranoside (BPMP) ligand as described previously (19, 43). *E. coli* and mammalian produced material bound BPMP ligand with similar affinity (**Fig. 1C**); thermal stability, as determined by differential scanning calorimetry, was also comparable (**Fig. 1D**).

### Design of a full-length donor strand complemented FimH (FimH-DSG)

To mimic the presentation of FimH on the assembled fimbrial tip (20), and understand the contribution of FimH_PD_ in ability to elicit anti-FimH antibodies with the ability to prevent bacterial binding to mannose, we considered production of a full length FimH antigen. Full length FimH cannot be produced in *E. coli* without the chaperone FimC or as part of a pilus (16, 19, 44, 45). In previous studies in humans and non-human primates, full length FimH in complex with FimC was used as an immunogen (23–25, 33, 34). Whether the FimH_PD_ is important for optimal immunogenicity is unknown, therefore the design of a full length FimH was explored to compare immunogenicity with that of FimH_LD_ alone. Sauer *et al* previously produced a stable, isolated FimH molecule by displacing FimC in FimCH with a FimG donor strand peptide (FimG residues 1-14) (46). This approach requires coexpression of FimH with the chaperone, FimC. In contrast, Barnhardt *et al* produced a stable, single chain FimG donor strand complemented FimH in *E. coli* using a 4-residue amino acid linker (DNKQ), enabling production of full length FimH without a chaperone (17). The single chain concept was explored further in the current study. Donor-strand complemented, full-length FimH proteins (FimH-DSG) were produced by attaching a 14-mer donor strand peptide from FimG to the C-terminus of the FimH_PD_ via a Glycine-Serine linker of various lengths, as well as the previously described DNKQ linker (17). Flexible polypeptide linkers consisting of Glycine and Serine are often used in construction of multidomain proteins as these flexible and hydrophilic spacer sequences prevent formation of secondary structure between protein domains, reducing the possibility that linkers will interfere with the folding or function of the target protein (47). Constructs were screened in mammalian cells and expression levels were similar, as assessed by SDS-PAGE (**Fig. S1**). FimH-DSG with a 7-residue Glycine-Serine linker (GGSSGGG) was selected based as optimal based on structural analysis. Non-native glycosylation in FimH_LD_ was prevented by incorporation of N7S and N70S mutations as described above and an additional mutation to Serine was also introduced at N228 in the FimH_PD_. Analysis by mass spectrometry confirmed that FimH-DSG retained a single glycosylation site at N235 in the exploratory antigen characterized in this study (data not shown). Note, fully aglycosyl variants of FimH-DSG were subsequently generated following introduction of an additional N235S mutation and expression in ExpiCHO cells (described in brief below).

The proposed mechanism of action of FimH-containing vaccines to prevent UTIs is via inhibition of bacterial binding to the urinary tract epithelium (25). We developed a live, whole *E. coli* binding inhibition assay to quantify the ability of anti-FimH antibodies to prevent binding of fimbriated bacteria to a mannosylated surface (mimicking the mannosylated FimH receptor on the surface of bladder epithelial cells), based on previous work (48). Using this assay, the relative ability of *E. coli* and mammalian derived FimH proteins to elicit inhibitory antibodies in mice was evaluated (**Fig. 1E** and **Fig. 1F**). Inhibitory titers elicited by *E. coli* and mammalian derived WT FimH_LD_ proteins in mice were equivalent, although the potency of the stabilized lock mutant FimH_LD_ V27C L34C produced in mammalian cells was slightly lower than that of the *E. coli* produced FimH_LD_ V27C L34C. Disulfide bond formation may differ in the periplasm of *E. coli* compared to the mammalian cytoplasm (49). Inhibitory titers elicited by full length FimH proteins (FimH-DSG, only produced in mammalian cells) including the previously described FimCH protein (50) were markedly higher than FimH_LD_ proteins in terms of percentage of responders and geometric mean IC_50_ values (**Fig. 1F** and **Table S1**).

### Rational design of FimH variants stabilizing the low-affinity, open conformation

We employed *in silico* analysis to identify novel variants predicted to stabilize FimH_LD_ in a low affinity conformation, which is unable to bind its cognate mannose receptor. Crystal structures of FimH in complex with fimbrial structural proteins in the absence of ligand or presence of D-mannose are shown in **Fig. 2A**, illustrating the differences in FimH_LD_ conformation. The crystal structure of full length FimH in complex with fimbrial structural proteins (PDB ID 3JWN (20)) was used as a model. Suggested amino acid substitutions identified from this analysis were introduced into either FimH_LD_ or FimH-DSG (**Fig. 2B**). To stabilize FimH_LD_ in an open conformation, the following design strategies were applied: 1) nonpolar residues exposed to solvent in the pre-bound state but buried within the protein interior in the bound structure were mutated to polar or charged residues, disfavoring the high-affinity, closed conformation of FimH_LD_, 2) disulfide linkages were introduced between residue pairs in close proximity in the pre-bound state, but not in the bound conformation, 3) mutation of glycine residues having a backbone F-angle < 0° in the pre-bound state but > 0° in the bound structure to prevent closure of the ligand binding site, 4) a full length, single chain FimH-DSG was designed (described above) 5) cavity filling mutations designed to stabilize the interface between FimH_PD_ and FimH_LD_ were introduced separately into full length FimH-DSG.

**Fig. 2.**
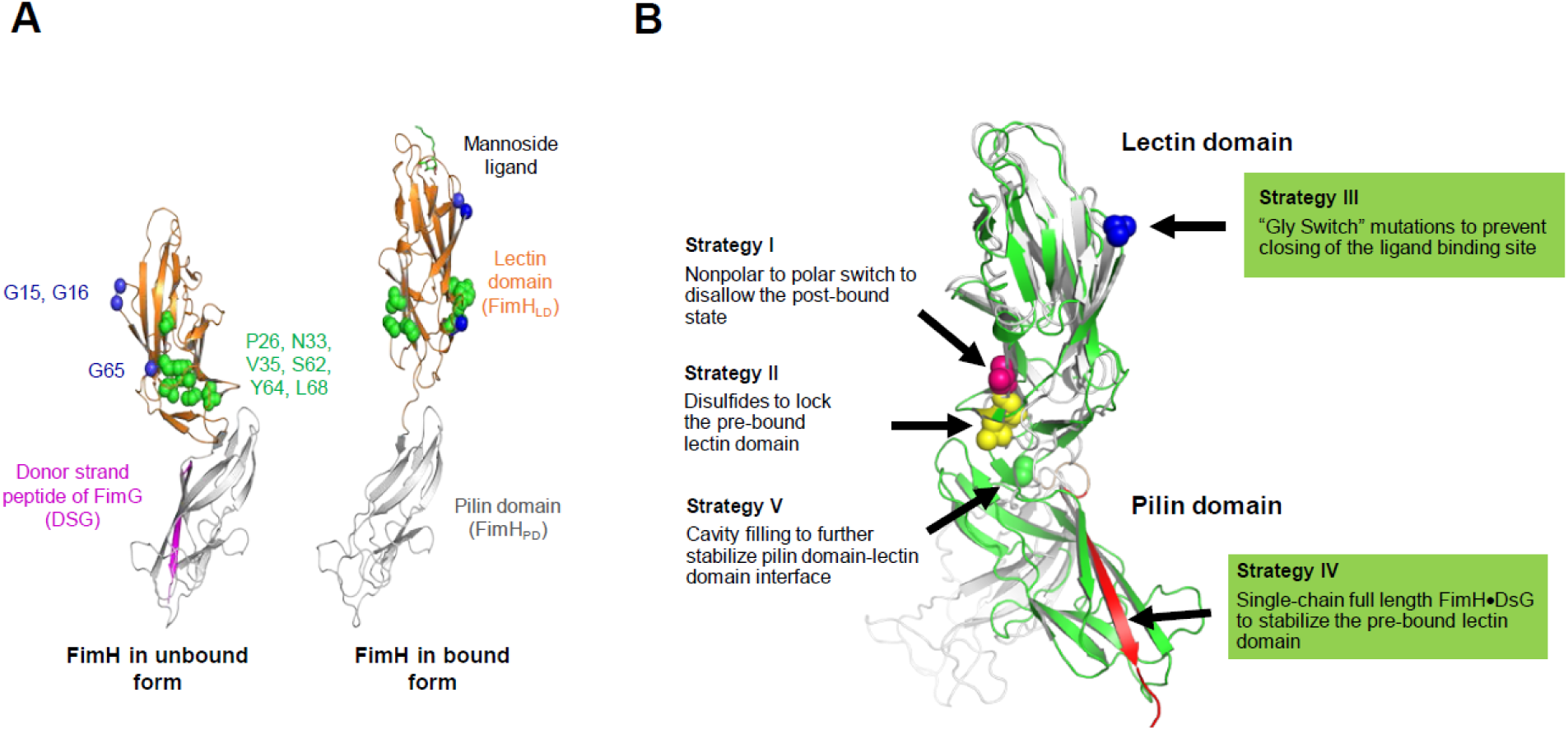
Rational design of FimH mutations stabilizing the native state. (**A**) Structures of FimH in unbound form (PDB structure 3JWN) and bound to mannoside ligand (PDB structure 1KLF). In pili, unbound FimH (left), complexed with the donor strand of FimG, adopts a compact conformation that binds the FimH cognate receptor, the terminal mannose moiety of glycosylated proteins, with low affinity. Upon binding a mannose moiety (right), the FimH_LD_ and FimH_PD_ separate, sidechains (colored in green) flip from protein interior to surface, and backbones of Gly residues (colored in blue) exhibit large conformational changes. Residues shown in blue and green were targeted for mutagenesis. **(B)** Strategies employed to stabilize FimH conformation. Unbound (grey) and bound (green) full length FimH structures with residues targeted by mutagenesis are highlighted.

### Screening of FimH_LD_ and FimH-DSG mutants in *in vitro* assays

Sixty-four mutant and WT versions of FimH_LD_ and FimH-DSG proteins were expressed in Expi293 cells as described above. Purified proteins were evaluated in a series of *in vitro* and *in vivo* studies (**Fig. 3A**). Mutants that were expressed at low levels were excluded from further evaluation. Binding affinities (K_d_) of FimH mutants for mannoside ligand were determined by FP assay using BPMP as described above. K_d_ values for a subset of FimH_LD_ and FimH-DSG mutants relative to WT are shown in **Fig. 3B** (data for additional mutants can be found in **Table S2**). The sequence of FimH_LD_ WT is derived from *E. coli* UTI isolate J96 (51). V27A is a natural variant that is associated with virulent UTI isolates and those associated with Crohn’s Disease (11, 52). Introduction of the single point mutant V27A did not alter mannoside ligand binding affinity relative to FimH_LD_ WT, while combining single or double glycine loop mutations at G15 or G16 positions with V27A significantly impaired ligand binding (**Table S2**). Ligand binding was completely lost in the triple mutant, FimH_LD_ G15A G16A V27A (FimH_LD_ TM, K_d_ > 2000 nM). Ligand binding affinity of FimH-DSG WT was reduced more than 100-fold compared to FimH_LD_ WT, in agreement with the allosteric role of the pilin domain in regulating ligand binding by the lectin-binding domain. FimH-DSG V27A also had a two-fold lower ligand-binding affinity, compared to FimH-DSG WT (**Table S2**). This is consistent with previous data showing that a FimH mutant containing A27 has the propensity to adopt a less-active state that binds mannose with low affinity (11). Like FimH_LD_, FimH-DSG mutants containing V27A and Glycine loop mutations have substantially reduced ligand binding activity. Altogether, our data suggest that the flexible Glycine loop plays a stabilizing role in ligand binding. Mutations in this loop result in FimH adhesins with poor affinity for the synthetic BPMP ligand used as representative of cognate mannosylated host cell glycoprotein receptors.

**Fig. 3.**
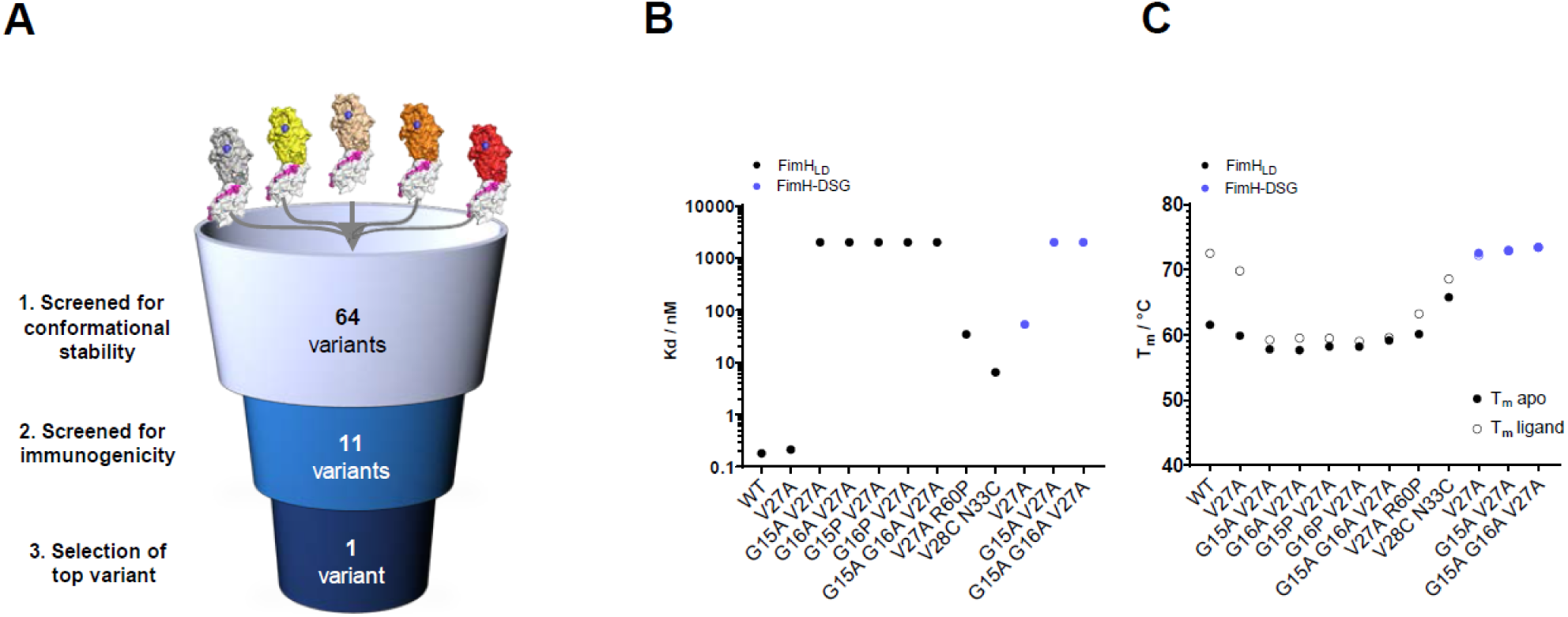
Identification of FimH mutants with improved thermal stability and reduced mannoside ligand affinity. **(A)** 64 FimH variants were screened *in vitro* for ability to bind mannose, thermal stability, and conformation. A subset of constructs was screened for immunogenicity in mice. **(B-C)** Biochemical characterization of purified FimH_LD_ and FimH-DSG mutants. **(B)** relative average binding affinities of FimH mutants to mannoside ligand. Note, the assay limit of detection was ∼2000 nM. **(C)** filled circles display average melting temperatures of each mutant. Open circles denote melting temperature of FimH protein in the presence of mannoside ligand. Tabulated K_d_ and T_m_ values for all mutants are in **Tables S2** and **S3** respectively.

To further evaluate ligand binding abilities of FimH variants, a SYPRO orange-based differential scanning fluorimetry assay was used (19). The temperature at which 50% of the protein is unfolded (T_m_) was determined for all mutants in apo state as well as in the presence of methyl alpha-D-mannopyranoside, a derivative of alpha-D-mannose that binds to FimH with micromolar affinity (53). The T_m_ of apo protein and its corresponding T_m_ shift (ΔT_m_) in ligand bound condition are summarized in **Table S3**. FimH-DSG WT proteins exhibited significantly higher T_m_ (71.7 °C) compared to FimH_LD_ WT (61.5 °C). T_m_ shifts of the FimH mutant proteins from apo to mannoside ligand bound conditions are in concordance with the changes seen in mannoside ligand K_d_ measurement (**Fig. 3B, Table S2**). Of note, consistent with published data the V27C L34C disulfide lock mutant (19) exhibited significantly lower thermostability (T_m_) compared to FimH_LD_ and FimH-DSG WT proteins, while retaining mannoside ligand binding activity (Kd of 30-60 nM). In contrast, combinatory mutations containing V27A and Glycine loop mutations at G15 and G16 positions resulted in FimH mutants that retained thermostability while losing their ability to bind mannoside ligand (Kd >2000nM). Together with our binding affinity data, these data suggest that combining V27A and Glycine loop mutations stabilizes FimH proteins in a low affinity state that is largely incapable of ligand binding.

In previous work, WT and conformationally locked FimH_LD_ mutants were found to have distinct tertiary structures (19). The secondary and tertiary structures of selected FimH_LD_ and FimH-DSG proteins were therefore examined by CD (**Fig. S2**). The far-UV CD spectrum (secondary structure) of FimH_LD_ WT expressed in mammalian cells is consistent with previously published data. Overall, the secondary structures of the FimH_LD_ or FimH-DSG mutants are highly similar to WT proteins (**Fig. S2A**), suggesting that the overall secondary structure is not altered in these mutants. The near-UV CD spectra (tertiary structure) of most FimH_LD_ mutants except for V27A (**Fig. S2B**), however, were quite different from that of FimH_LD_ WT and are more similar to the previously reported FimH_LD_ V27C L34C and V27A R60P mutants which assume an open conformational state (19). On the other hand, all FimH-DSG proteins including WT and the V27A, G15A, G16A triple mutant (or TM) had highly similar secondary and tertiary structures, resembling the open state of FimH_LD_ (**Fig. S2**). Overall, the CD characterization is consistent with mannoside ligand binding data described above, which suggests these mutants are stabilized in an open conformation that has low mannose binding affinity.

Together, these data suggest that the Glycine loop mutations constrain FimH_LD_ in an open conformation, while the FimH-DSG constructs including the corresponding reference WT variant, remain in a conformation that remains unchanged upon introduction of stabilizing mutations that eliminate ligand binding.

### FimH-DSG mutants induce antibodies with superior ability to inhibit bacterial binding than FimH_LD_ mutants in mice

To evaluate the relative ability of selected FimH mutants to elicit inhibitory antibodies, mice were vaccinated with each mutant protein and the collected sera were used to assess the ability to prevent bacterial binding *in vitro.* Mice were immunized three times with 10 µg FimH protein combined with QS21 adjuvant (**Fig. 4A**) and sera from post dose 2 and 3 time points were evaluated in the yeast mannan *E. coli* binding inhibition assay described above (**Fig. 4B** and **Table S6**).

**Fig. 4.**
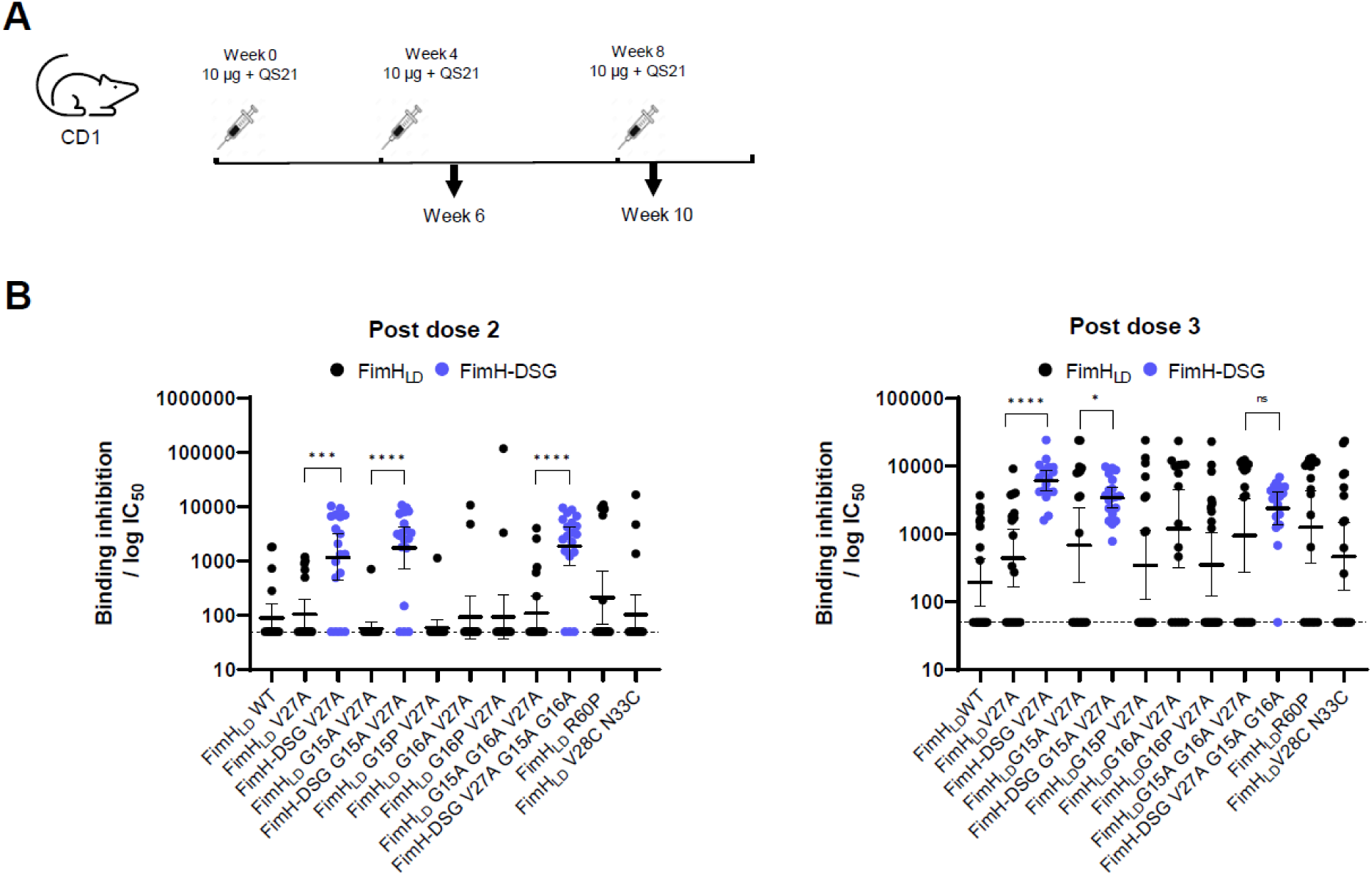
FimH-DSG mutants induce antibodies with superior ability to inhibit bacterial binding compared to FimH_LD_ mutants in mice. **(A)** CD-1 mice were immunized 3 times with 10 µg FimH with QS21 adjuvant. Sera were analyzed for the ability to block FimH-expressing *E. coli* binding to yeast mannan. **(B)** Inhibitory titers were determined from serial dilution of sera from vaccinated mice and represent the reciprocal of the dilution of serum at which 50% of bacteria remain bound to the plate and are shown for post dose 2 and post dose 3 timepoints. Statistical significance (*p-*value) of differences in responses between groups was determined using an unpaired t-test with Welch’s correction applied to log-transformed data; the bars and asterisk illustrate the significance of the difference in response between groups. Tabulated IC_50_ values are shown in supplemental **Table S6**.

At post dose 3, mice immunized with novel stabilized FimH_LD_ mutants G15A G16A V27A, G16A V27A, and the previously described mutant V27A R60P (19) had higher responder rates and increased binding inhibition (*p* <0.05) compared to FimH_LD_ WT (**Fig. 4B** and **Table S6**). Other FimH_LD_ mutants (G15A V27A, G16P V27A, V28C N33C) did not significantly enhance functional immunogenicity of FimH_LD_. Thus, several mutants designed to enhance functional immunogenicity of FimH_LD_ by constraining FimH_LD_ in an open conformation led to improved responses relative to FimH_LD_ WT, confirming previous findings (54). Following vaccination with 2 doses of FimH_LD_ and FimH-DSG proteins, significantly more animals yielded inhibitory titers in the groups vaccinated with FimH-DSG compared to FimH_LD_ (**Fig. 4B** and **Table S6**). This trend was sustained at PD3, where 95%-100% of mice responded in groups vaccinated with FimH-DSG V27A, FimH-DSG G15A V27A or FimH-DSG TM and tended to have higher GMC IC_50_ values relative to FimH_LD_ mutants (*p*<0.05) (**Fig. 4B** and **Table S6**). In conclusion, FimH-DSG mutants elicit higher inhibitory antibody responses compared to FimH_LD_ mutants.

The ability of FimH-DSG V27A, G15A V27A and the G15A G16A V27A triple mutant (TM) to elicit inhibitory antibodies were similar. However, during large scale purification of FimH-DSG WT and TM from ExpiCHO cells for further characterization, a tendency for aggregation was observed for FimH-DSG WT and characterization by analytical size exclusion chromatography revealed the presence of high molecular weight complexes (**Fig. S3A-C**). We hypothesized that during CHO fermentation and upon secretion into the culture media the FimH-DSG WT binds glycan molecule(s) released from the surface of the host CHO cells. To evaluate this further, samples were analyzed by High pH Anion-Exchange Chromatography with Pulsed Amperometric (electrochemical) Detection (HPAEC-PAD). FimH-DSG WT preparations contained numerous monosaccharides (**Fig. S3D**). In contrast, FimH-DSG TM was bound to comparatively fewer glycan moieties (**Fig. S3D**). Furthermore, it is entirely possible that the low monosaccharide content that was detected represents sugar moieties of the N-glycan present on N235. FimH-DSG TM did not tend to aggregate or form high molecular weight complexes. Therefore, as it can be purified to homogeneity, at a high yield, we used this mutant for our subsequent studies.

### Production of aglycosylated FimH-DSG TM

Removal of the N-glycan present on residue N235 of FimH-DSG TM led to additional glycosylation on residue N228. To avoid these issues, we explored removal of glycosylation by introducing the following combinations of mutations: N228S N235S, T230A T237A, N228G N235G, and N228Q N235Q. Proteins were expressed transiently in 2L ExpiCHO cell cultures and yields were between 69 mg (N228G N235G) and 311 mg (N228S N235S). Aglycosylation mutations did not impact the ability of inhibitory Mabs to bind relative to glycosylated FimH-DSG TM (**Table S7**), and the capability of aglycosylated variants elicit adhesin blocking antibodies does not appear to differ from the glycosylated parent (**Fig. S4**).

### Structural characterization of FimH-DSG TM by X-ray Crystallography

Among all conformationally stabilized FimH-DSG mutants, FimH-DSG TM was of most interest as it harbors the two Glycine loop mutations G15A and G16A designed to conformationally lock the mannose site in the ‘open’ state. To verify the conformation of FimH-DSG TM, the crystal structure of a FimH-DSG TM protein containing four glycosylation site mutations (N7S, N70S, N78S, and N228Q) was solved by X-ray crystallography at 1.9 Å resolution (**Fig. 5A** and **Table S9**). There are four molecules of FimH-DSG TM in one asymmetric unit which share high similarity, therefore, structural analysis was done with protomer A. Introduction of the glycosylation site mutations did not appear to alter either the local or global conformations of the FimH_LD_ and FimH_PD_. Comparison of the structure to previously determined structures of FimH_LD_ confirmed that FimH-DSG TM lectin domain adopts a compressed structure with an open mannose-binding pocket (**Fig. 5C**). Clear electron density was obtained for the Glycine loop, wherein the G15A G16A mutations were introduced which induced a local change in the loop conformation (**Figs. 5B, 5C**). A15 projects towards the inside region of the clamp loop, providing rigidity, which causes the loop to widen at the end by ∼1.2 Å (Ile13 Ca-Ser17 Ca) compared to the WT residue (G15), thus stabilizing the low affinity state. Additional ridigity is introduced by the G16A substitution, as it projects towards the bulk solvent. By comparison, these residues adopt very different conformations in the closed, high-affinity state (55). The overall structure of FimH-DSG TM resembles the low affinity conformation of FimH_LD_ (**Fig. 5D**) (16, 18). In comparison to previously published structures, the donor strand complemented FimH_PD_ did not exhibit any conformational changes. The backbone of the exogeneous 7-residue Glycine-Serine linker that connects the FimH and FimG peptide could largely be resolved, and does not impact on the conformation of the lectin or pilin domains in comparison to previously published structures (**Fig. S5**) (18).

**Fig. 5.**
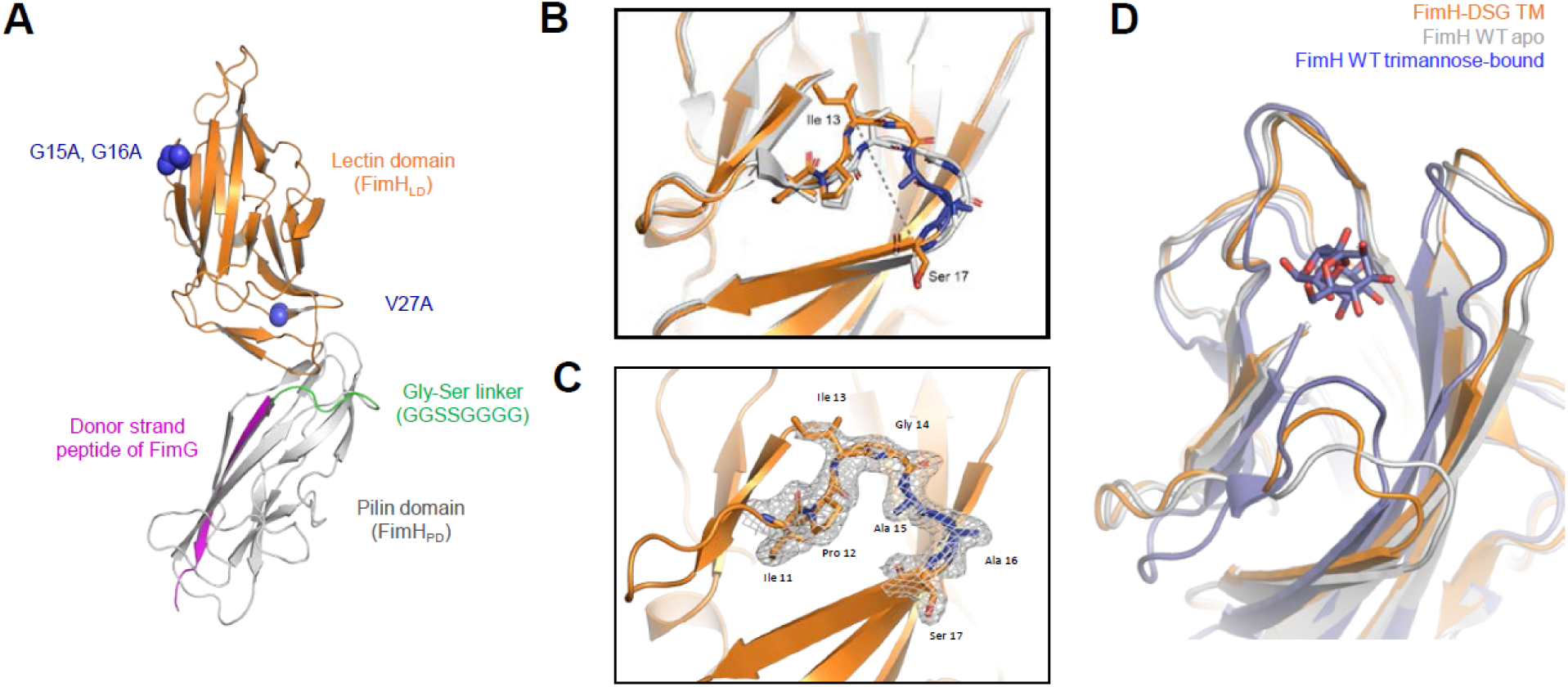
The lectin domain of FimH-DSG TM adopts an open conformation. **(A)** Overall structure of FimH-DSG TM from x-ray crystallography data (PDB code 8V3J). **(B)** Superimposition of ligand binding sites of FimH-DSG TM (orange) and WT FimH from a previously published structure of native pili (PDB code 3JWN, light grey) shows remodeling of the Glycine loop due to G15A G16A mutations. The widening of the loop between Ile13 Ca-Ser17 Ca is shown in a dotted line. **(C)** Electron density and atomic model of Glycine loop in FimH-DSG TM. **(D)** Superimposition of FimH-DSG TM (orange), WT FimH in apo (PDB code 3JWN, light grey) and trimannose-bound (PDB code 6GTV, slate) forms shows that the ligand binding site of FimH-DSG TM adopts an open conformation resembling the apo state but not the ligand-bound, closed conformation.

### Inhibitory antibodies elicited by FimH-DSG map to novel epitopes

As described above, FimH-DSG WT and derived mutants induced superior inhibitory antibody responses in mice compared to equivalent FimH_LD_ variants. To further characterize inhibitory antibodies induced by FimH-DSG, monoclonal anti-FimH antibodies were developed and screened by the yeast mannan *E. coli* binding inhibition assay, which resulted in the identification of 10 unique clones (**Table 1**).

**Table 1.** Identification of novel FimH inhibitory antibodies. Mabs against FimH were screened for ability to inhibit bacterial binding in the bacterial inhibition assay. Resulting Mabs were subjected to a series of competition experiments to identify groups of antibodies likely binding similar epitopes. This table shows inhibitory titers of each antibody and relative FimH binding affinity. Two Mabs, 440-2 and 445-3, were derived from mice immunized with a conformationally locked version of FimH, FimH-DSG LM (V27C L34C).

|  |  |  | Inhibitory<br>(IC<sub>50</sub>) titer,<br>μg/ml | Kinetic Measurements with FimH-DSG WT (Averages with Stdev) |  |  |  |  |  |  |
| --- | --- | --- | --- | --- | --- | --- | --- | --- | --- | --- |
| Site | Mab type | FimH<br>Mabs |  | k<sub>a</sub> (1/Ms) | k<sub>a</sub><br>(1/Ms)<br>Stdev | k<sub>dis</sub> (1/s) | k<sub>dis</sub><br>(1/s)<br>Stdev | K<sub>D</sub><br>(nM) | n | Elicited<br>antigen |
| 1 | Reference | 475 | 26.4 | 1.9E+05 | ±2.6E4 | 5.7E-04 | 1.8E-04 | 3.1 | 8 |  |
|  |  | 926 | 0.4 | 1.9E+05 | ±5.3E4 | 6.5E-04 | 4.8E-05 | 0.9 | 5 |  |
| 1 | Ligand<br>Binding<br>epitope | 299-3 | 1.4 | 4.1E+05 | ±2.5E4 | 3.4E-04 | 3.3E-05 | 0.8 | 4 | FimH-DSG WT |
|  |  | 304-1 | 0.3 | 4.1E+05 | ±4.3E4 | 1.0E-04 | 1.6E-05 | 0.2 | 7 |  |
|  |  | 306-2 | 4.9 | 4.1E+05 | ±3.0E4 | 4.8E-04 | 1.5E-05 | 0.6 | 5 |  |
|  |  | 313-1 | 11.9 | 4.1E+05 | ±2.3E4 | 5.3E-04 | 1.2E-05 | 2.1 | 5 |  |
|  |  | 330-2 | 14.3 | 4.1E+05 | ±8.1E4 | 1.1E-03 | 1.6E-05 | 1.9 | 5 |  |
|  |  | 338-4 | 2.5 | 4.1E+05 | ±9.7E3 | 2.1E-04 | 1.4E-05 | 1.4 | 5 |  |
| 2 | Non-<br>ligand<br>binding<br>site<br>epitope | 327-3 | 1.0 | 5.8E+05 | ±1.5E4 | 8.0E-05 | 6.3E-05 | 0.1 | 5 |  |
|  |  | 329-2 | 3.3 | 5.8E+05 | ±3.5E4 | 6.7E-05 | 1.8E-05 | 0.1 | 5 |  |
| 3 | Non-<br>ligand<br>binding<br>site<br>epitope | 445-3 | 1.3 | 5.8E+05 | ±4.2E4 | 7.5E-05 | 2.8E-05 | 0.1 | 5 | FimH<br>DSG-LM |
| 4 | Non-<br>ligand<br>binding<br>site<br>epitope | 440-2 | 94.8 | 9.4E+04 | ±1.9E4 | 6.2E-04 | 8.2E-05 | 7.0 | 5 |  |

A series of competition experiments using Bio-layer interferometry (BLI) was undertaken to facilitate Mab classification and epitope mapping. As comparators, two previously described inhibitory Mabs, 475 and 926, which bind to overlapping epitopes on the ligand binding interface, were included (48, 54). Experiments were performed with and without octylmannopyranoside ligand, to assess whether there is competition between the ligand and Mab binding to FimH_LD_ (**Table 1, Fig. S6)**. These experiments identified four distinct inhibitory epitopes. Site 1 was recognized by 6 novel Mabs, as well as reference Mabs 926 and 475, which effectively block ligand binding. Three Mabs were unable to block ligand binding and bound to two distinct epitopes (site 2 and site 3, composed of 327-3 and 329-2, and 445-3 respectively). Site 4 was recognized by a single Mab (440–2) that bound exclusively to conformation-stabilized FimH_LD_. Note, this Mab was derived from mice immunized with the conformationally locked construct, FimH-DSG V27C L34C. Individual Mab binding kinetics are shown in **Table 1** alongside their inhibitory concentrations in the *E. coli* binding inhibition assay.

Fab fragments derived from representative antibodies from each new site (329-2 (site 2), 445-3 (site 3) and 440-2 (site 4)) were selected as representatives for subsequent epitope mapping by cryo-EM. The small size of FimH was an impediment to high resolution structural analysis of FimH-Fab binary complexes, which yielded only low-resolution structures that could not be modeled. To circumvent this limitation, a combinatorial approach was taken in which pairs of Fabs with non-overlapping epitopes were combined with FimH to form ternary complexes, increasing the overall particle mass to 125 kDa. Using this ternary complex strategy, a cryo-EM structure of FimH-DSG TM with Fabs 329-2 and 445-3 was determined at 3.11 Å (**Fig. 6A, Fig. S7, Table S10**), with the final map comprising FimH_LD_ and the antigen-binding portions of each Fab, allowing molecular modeling of each component and identification of the molecular details of the respective epitopes. The two Fabs were distinguished in part by comparison of their sequences to the side chains in the map, but to build additional confidence, the structure of a similar FimH-DSG TM ternary complex, in which the 329-2 Fab was replaced with 440-2, was determined by cryo-EM (**Fig. 6B, Fig. S8, Table S10**). While the new map from a 3.9 Å reconstruction could be used to identify density corresponding to FimH and both Fabs, the map resolution and quality were limited by under-represented particle views, allows only rigid body fitting of models. Comparison of the Fab positions in both structures, however, confirmed the Fab assignments for 329-2 and 445-3, and further allowed mapping of the 440-2 epitope. Surprisingly, Mab 440-2 binds to the tip of FimH_LD_, which covers the mannose binding site and largely overlaps with the known site 1. Binding of Mab 440-2 (site 4) therefore directly competes with mannose binding, conveying an orthosteric inhibition mechanism for blocking of bacterial adhesion, like that observed for the competitive Mab 475 (48). However, being derived from a conformationally locked FimH-DSG mutant distinguishes Mab 440-2 from other inhibitory Mabs. Mab 440-2 preferentially binds to the low affinity open state of FimH_LD_, but not WT FimH_LD_ (**Table S4**), suggesting that it uses a novel inhibitory mechanism. Indeed, the structure of the ligand binding site of 440-2-bound FimH_LD_ matches the open conformation, largely distinct from the observed conformation in the ligand bound high-affinity ‘closed’ state (**Fig. 7A**).

**Fig. 6.**
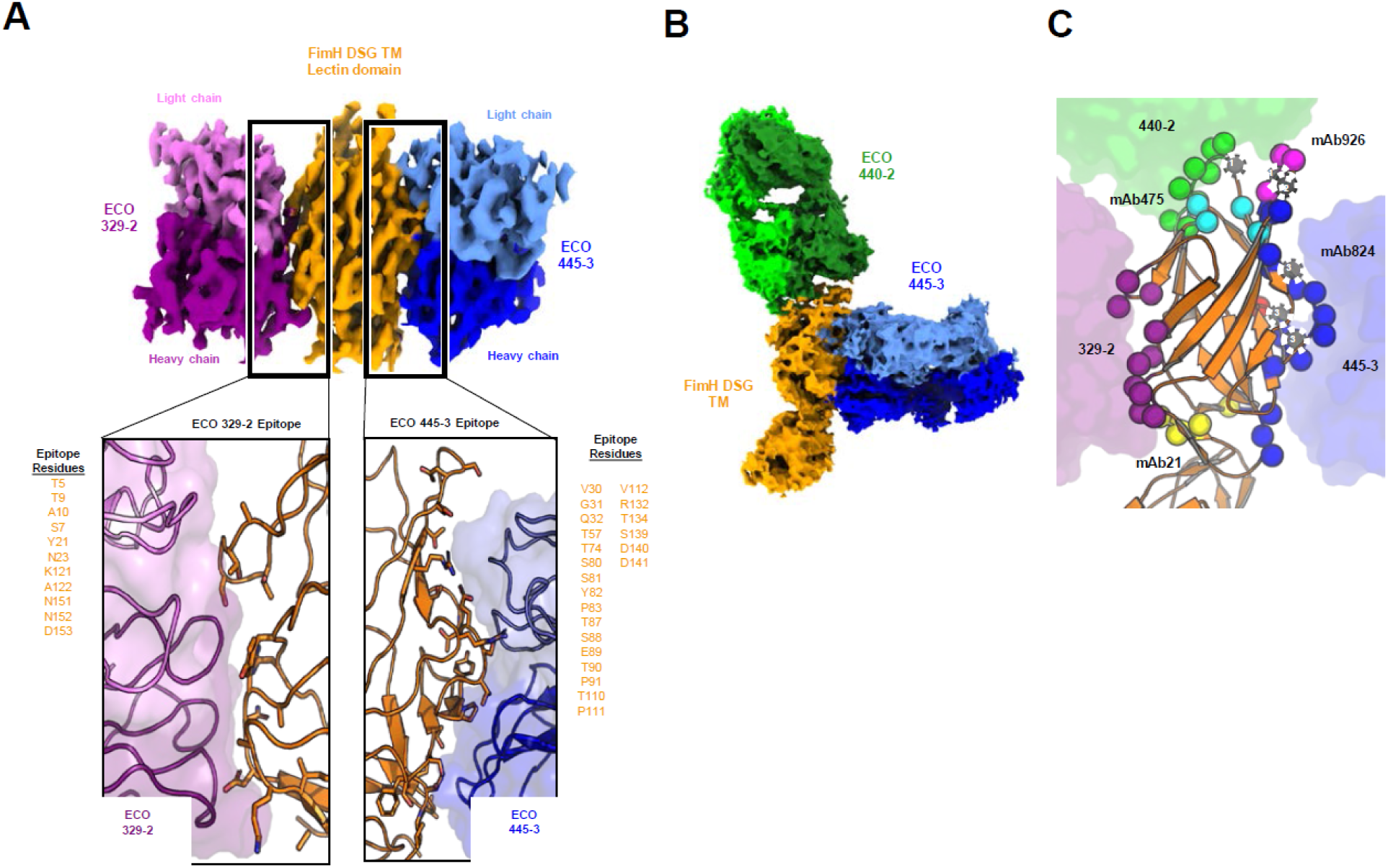
Identification of novel inhibitory epitopes on FimH-DSG TM. **(A)** Structure of FimH-DSG TM in complex with 329-2 and 445-3 Fabs solved by cryo-EM. Top image shows the cryo-EM map, colored by chain, as indicated. Insets show respective epitope interfaces, with FimH residues contributing to each epitope surface shown in stick representation and listed along the side. **(B)** CryoEM structure of FimH-DSG TM in complex with 440-2 and 445-3 Fabs, colored by chain, as indicated. While this map was not suitable for modeling, the Fabs and respective epitopes could be distinguished by comparison to the structure in panel A. **(C)** Cartoon representation of the FimH_LD_ in the FimH-DSG TM crystal structure. Epitopes identified in this study and in previous work by others (Mab 926, 824, 475 and 21) are highlighted, with participating residues shown as spheres and colored as indicated. Numbered gray residues indicate coincidence between two epitopes (1 = Mab 926 and Mab 475, 2 = Mab 926 and Mab 445-3, 3 = Mab 475 and Mab 824).

**Fig. 7.**
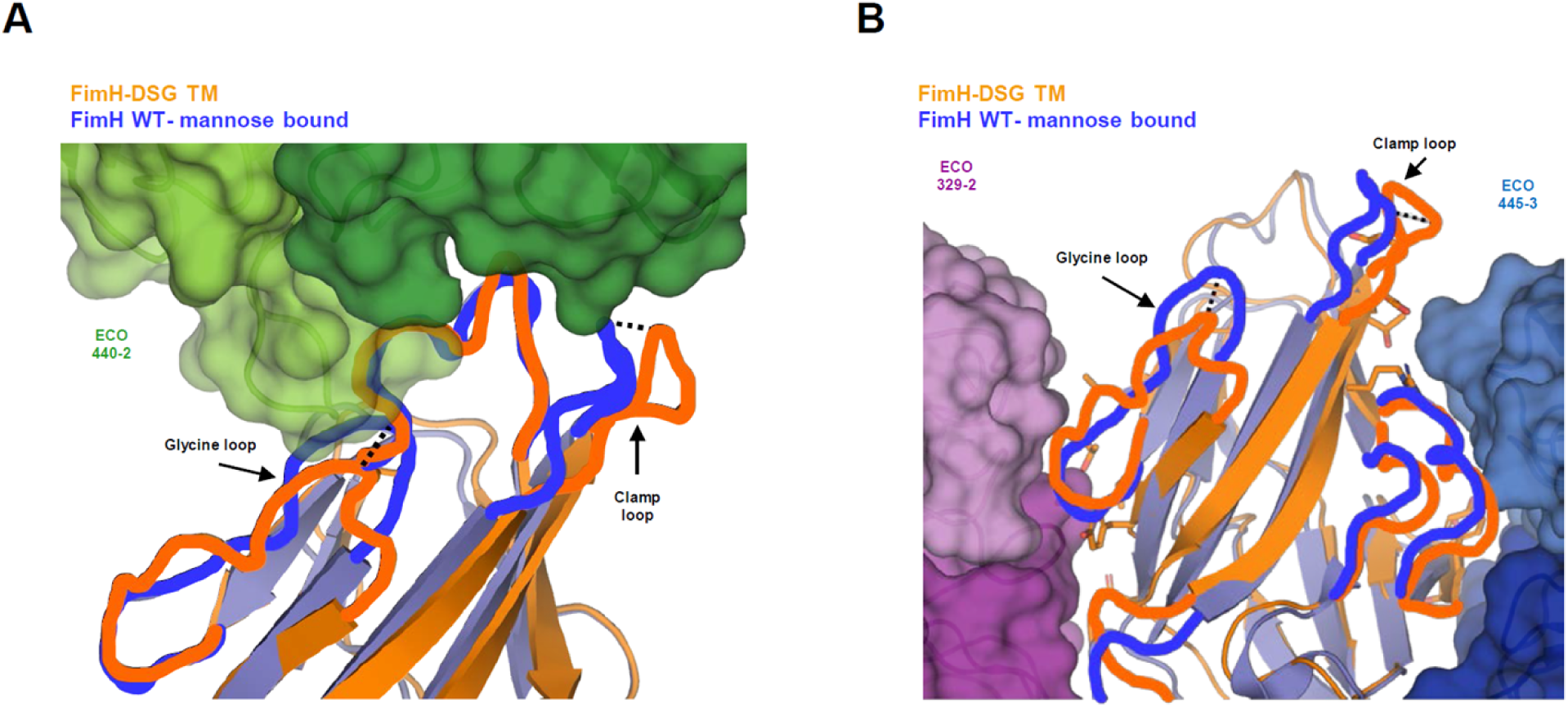
Structural basis of inhibitory mechanisms of different Mabs. **(B)** Superimposition of FimH WT bound to trimannose (PDB code 6GTV, blue) and FimH-DSG TM (orange) cryo-EM structure resolved from the complex with Fabs of 440-2 (green surface) and 445-3 (not shown). **(B)** Superimposition of FimH WT bound to trimannose (PDB code 6GTV, blue) and FimH-DSG TM (orange) cryo-EM structure resolved from the complex with Fabs of 329-2 (purple) and 445-3 (dark blue). Surface representations for the two Fabs are shown. Side chains from FimH-DSG TM that are part of the Mab epitopes are shown in stick format. In **(A)** and **(B)**, shift of the clamp loop and Glycine loop from FimH-DSG TM to mannose-bound FimH WT is highlighted by dashed lines. Conformations of the important FimH epitope loops for 440-2, 329-2, and 440-2 binding are highlighted by orange (FimH-DSG TM structure) and blue (FimH WT-mannose bound structure) lines.

External to the ligand binding site, epitopes of Mabs 329-2 (site 2) and 445-3 (site 3) were mapped to the opposite sides of FimH_LD_. Despite not overlapping with ligand binding pocket, the 329-2 and 445-3 epitopes are composed of residues from the flexible loops that are part of the ligand binding site, with conformations that change significantly during the switch between inactive and active states. In particular, Mab 329-2 recognizes a novel inhibitory epitope comprised of residues T5, S7, T9, A10, Y21 and N23. These residues are connected to the Glycine loop that harbors the G15A and G16A mutations. As a result, binding of Mab 329-2 stabilizes the Glycine loop in the open conformation that has low mannose affinity (**Fig. 7B**). Binding to the other side of FimH_LD_, Mab 445-3 interacts with residues including R132, T134, S139, D140 and D141 from the clamp loop in its open conformation, which shifts 4.4 Å away from the compact-mannose-bound conformation seen in the closed form (PDB ID 6GTV, **Fig. 6B**). In addition, S80, S81, Y82 and P91 were also mapped to 445-3 binding site. These residues were revealed in previous work as being part of a critical epitope for Mab 824, an inhibitory antibody that allosterically prevents transitions of FimH from low-affinity to high-affinity states (56). Taken together, Mab 445-3 likely blocks bacterial adhesion via a combinatory allosteric mechanism which modulates both the global state transition as well as the local ligand binding site. These new epitopes described herein, along with previously identified FimH Mab epitopes are highlighted on the structure of FimH-DSG TM in **Fig. 6C**.

Of note, none of the epitopes mapped for these three Fabs overlap with the conformation stabilizing mutations introduced into the FimH-DSG TM vaccine candidate. Collectively, these results broaden the repertoire of FimH_LD_ functional epitopes, highlighting the diverse interference mechanisms by which antibodies contribute to FimH vaccine efficacy.

## Discussion

FimH is a key target for UTI vaccines due to its integral role in UPEC virulence. However, clinical trials have yet to demonstrate a clear protective effect of FimH immunization in humans (33). As there are issues with scalable production and poor immunogenicity documented in previous studies, we sought to optimize the antigen design and how it is bioprocessed to improve its application for therapeutic use.

Firstly, we developed a mammalian expression system to produce FimH antigens. In contrast to classic osmotic shock approaches that release functional FimH from the *E. coli* periplasm, the mammalian expression platform dramatically increases yield and simplifies the purification process. Notably, FimH antigens produced in mammalian cells are structurally and antigenically alike to the analogous proteins produced in *E. coli*. Our approach provides a path forward for manufacture of FimH antigens in large quantities for clinical trials and may be applicable to other complementary fimbrial vaccine antigens, that are similarly expressed at low levels in *E. coli*.

Computational design has been used successfully to optimize vaccine antigens for improved stability, immunogenicity (57, 58) as well as safety (57, 59, 60). The design of conformationally stabilized proteins to improve neutralizing responses was pioneered in the field of respiratory syncytial virus (RSV) research (38) and has been employed for COVID-19 (39), HIV (40), influenza (41), and malaria vaccines (61). In agreement with previous reports (54) our rational approach to engineering structurally constrained FimH mutants led to identification of an conformation-stabilized full-length FimH variant (FimH-DSG TM) that elicits superior bacterial binding inhibition when compared to WT or conformation-stabilized mutants of FimH_LD_. FimH_PD_ is thought to act as an allosteric inhibitor of the FimH_LD_ and interaction of FimH_LD_ with FimH_PD_ in full length FimH stabilizes the FimH_LD_ in the low-affinity state (62). Similarly, full length FimH-DSG TM is stabilized in the low affinity state (**Fig. 5**). Introduction of conformation stabilizing mutations (*e.g.,* G15A, G16A, V27C L34C) into FimH-DSG did not dramatically impact binding affinity nor immunogenicity further. As no inhibitory Mabs were found to bind FimH_PD_ in screening of FimH Mabs, the FimH_PD_ thus likely contributes to the increased immunogenicity of FimH-DSG compared to FimH_LD_ mutants via stabilization of the FimH_LD_ conformation. We hypothesize that this effect is due to exposure of binding pocket epitopes in the open conformation, enabling targeted antibody development against prebound FimH. Like other allosteric proteins, FimH can adopt different conformations, which, in turn, can affect accessibility and structure of functional (inhibitory) epitopes. Thus, different FimH_LD_ conformations may be antigenically distinct, and induce immune responses that differ in specificity or inhibitory function. This is well-documented for several viral proteins, including RSV fusion protein F: immunization with the prefusion-stabilized conformer induces a more potent neutralizing antibody responses, unlike the responses induced by the postfusion form(38). Similar observations have been made for SARS-CoV-2 Spike protein (63) and HIV-1 envelope protein (64).

Characterization of inhibitory Mabs elicited by FimH-DSG by epitope binning and cryo-EM revealed three non-overlapping epitopes present on FimH_LD_; indicating that inhibitory antibodies targeting FimH-DSG TM can act via more than one mechanism. Based on the characterization of antibodies raised against FimH_LD_, three types of FimH inhibitory antibodies are described in the literature (48, 62): orthosteric (where an antibody replaces ligand in the binding site), parasteric (where an antibody binds adjacent to ligand in the binding site), or dynasteric (where the antibody binds at an allosteric site distant to the ligand binding site, preventing a conformational switch). The analysis described herein mapped antibodies to the ligand binding site (site 1), as described previously, and three novel sites (sites 2, 3 and 4) that represent alternative novel allosteric mechanisms by which FimH directed antibodies can inhibit binding.

In addition to the improvements in immunogenicity and induction of antibodies against novel inhibitory epitopes, the single chain FimH-DSG TM antigen presented herein offers several advantages over existing FimH protein vaccine candidates. Production of FimH-DSG in mammalian cells yielded high levels of protein, which induced potent inhibitory titers in sera from immunized mice. FimH-DSG mimics the natural presentation of FimH in the context of the assembled pilus, without the need for a chaperone, resulting in a targeted immune response. Introduction of the three stabilizing mutations (G15A G16A V27A) enabled purification to homogeneity without copurification of contaminating glycans compared to WT FimH-DSG. The ability to express a fimbrial adhesin in mammalian cells at high yield, to homogeneity, in a highly immunogenic form, represents a key advancement in the production of fimbrial antigens for effective vaccine development.

## Conclusion

UTIs are a common problem and a primary source of sepsis in the immunocompromised and elderly (65). A vaccine which can decrease UTI may ultimately decrease rates of hospitalization and sepsis-associated morbidity and mortality. The optimized, highly immunogenic full length FimH antigen described in this study can be produced at scale in mammalian cells and induces potent inhibitory antibodies that have novel mechanisms of action. Thus, FimH-DSG TM is a promising candidate to evaluate further in efficacy studies against FimH-expressing *E. coli*.

## Materials and Methods

### Experimental Design

This work aimed to develop a *E. coli* vaccine candidate by applying a computational and experimental design strategy to enhance the stability of the FimH lectin binding domain (FimH_LD_) and single chain full length FimH (FimH-DSG). Mutations were designed to introduce amino acid substitutions in FimH_LD_ to stabilize the protein conformation in a low affinity state. FimH_LD_ and FimH-DSG mutants were expressed in mammalian cells and purified mutant proteins were characterized by thermal stability and rate of dissociation, in comparison to wild type FimH_LD_. The crystal structure of FimH-DSG G15A G16A V27A (FimH-DSG TM) was resolved to confirm the mutant’s open conformation. Monoclonal antibodies raised against FimH-DSG were evaluated in competition experiments to group into non-overlapping epitope bins. Representatives of each distinct bin were complexed with FimH-DSG TM and analyzed by Cryo-EM to determine epitopes.

### Ethics statement

Mouse immunogenicity studies were performed at Pfizer, Pearl River, NY, which is accredited by the Association for Assessment and Accreditation of Laboratory Animal Care (AAALAC). All procedures performed on mice were in accordance with local regulations and established guidelines and were reviewed and approved by an Institutional Animal Care and Use Committee (IACUC). The work was in accordance with United States Department of Agriculture Animal Welfare Act and Regulations and the NIH Guidelines for Research Involving Recombinant DNA Molecules, and Biosafety in Microbiological and Biomedical Laboratories.

### Expression and purification of WT FimH_LD_ from *E. coli*

DNA encoding J96 FimH_LD_ sequence was cloned into a pET28 vector and *E. coli* BL21(DE3) cells were transformed with the resulting construct. Expression, extraction and purification of FimH_LD_ or FimCH from the *E. coli* periplasm was performed as previously described (50, 66).

### Expression and purification of WT FimH and mutants in mammalian cells

DNA encoding J96 FimH sequence and its mutants was codon optimized for mammalian cells and cloned in frame behind sequence encoding mouse IgGκ or native FimH signal peptide, along with a C-terminal 8xHis tag in pcDNA3.1(+). FimCH was expressed from a pBudCE4.1 plasmid containing a CMV promoter to drive expression of FimC with a C-terminal 8xHis tag and a second promoter, EF1α, to drive expression of untagged FimH. For biochemical assays and mouse immunogenicity studies, endotoxin-free DNA was transiently transfected into Expi293 cells (ThermoFisher Scientific) according to manufacturer’s instructions. Supernatants were filtered through a 0.22 µm filter unit (Nalgene sterile disposable filter with PES membranes, ThermoFisher Scientific). Nickel Sepharose excel (Cytiva) was incubated with supernatants overnight at 4°C and purification was performed according to manufacturer’s instructions. The eluate was loaded onto a S200 16/600 column in 50 mM TrisHCl pH 8.0, 300 mM NaCl. Fractions containing pure protein were pooled.

For X-ray crystallography and cryoEM, FimH-DSG TM was produced in ExpiCHO cells (Thermo Fisher Scientific) according to manufacturer’s instructions. The purification procedure is described in the supplemental material.

### Computational design of FimH mutants stabilizing the native state

The crystal structure of native full length FimH in complex with fimbrial structural proteins (PDB ID 3JWN) was used as a model for the low-affinity, open conformation. The crystal structure of FimH_LD_ in complex with butyl α-D-mannoside was used as a model for the high-affinity, closed conformation (PDB 1UWF). Schrodinger BioLuminate (release 2017-2) was applied to analyze structural differences between the two states and to identify residue locations for mutations. Nonpolar residues that are exposed to solvent in the pre-bound structure and buried in the bound state were changed to polar or charged residues. Residue pairs selected for mutations to cysteine to form disulfide bonds that stabilize the native state were proximate (C_β_-C_β_ ≤ 5 Å) in the pre-bound state and distant (C_β_-C_β_ ³ 10 Å) in the bound conformation. Gly residues that have a negative backbone F-angle in the pre-bound state and a positive backbone F-angle in the bound state were mutated to Ala or Pro. An optimal Gly-Ser linker to connect the C-terminus of FimH and the N-terminus of FimG was identified using the Linker Modeler implemented in Molecular Operating Environment (MOE v2018, Chemical Computing Group).

### Characterization of WT and mutant FimH proteins

WT and mutant FimH proteins were characterized by fluorescence polarization, circular dichroism and differential scanning fluorimetry modified from Rabbani et al (19) and are described in the supplemental materials.

### Antigenicity assay

Inhibitory Mabs 299-3, 304-1 and 440-2 (developed in-house) were used to confirm the conformational state of FimH mutants; 299-3 and 304-1 bind to similar epitopes as Mab 475 and 926 (37, 48) while 440-2 recognizes a different epitope and appears to preferentially bind FimH_LD_ in an open conformational state. Mutants that maintain the same structure as WT FimH_LD_ are expected to bind all antibodies. Octet HTX from ForteBio was used for all the kinetic real-time biomolecular interaction experiments to measure antibody reactivity with each mutant. Experiments were carried out at 30 °C with 1000 rpm agitation in 96-well black plates containing 240 µl per well. Ni-NTA biosensors were equilibrated in buffer containing 1x PBS buffer containing 0.5 % BSA and 0.05 % Tween 20 (PBT) before allowing them to load his-tagged FimH mutant proteins at 5 µg/ml for 5 min. FimH loaded biosensors were allowed to reestablish baseline in PBT for 3 min before allowing them to associate with antibodies from different bins at 5 µg/ml for 5 min. Octet data analysis software was used for kinetic analysis of association step and obtain response in nm shift (tabulated).

### FimH whole cell binding inhibition assays

CFT073 (ATCC) was serially passaged in 10 ml of Luria Bertani broth in static growth conditions at 37°C to enrich for FimH expression. Surface expression reaching ≥95 % was confirmed via flow cytometry using anti-FimH Mab 926 (48). Prior to the assay, 384 well white Maxisorp plates (Nunc) were coated with 20 µg/ml of yeast mannan (Sigma-Aldrich) and blocked in 1 % BSA (Thermo). FimH-expressing *E. coli* cells (confirmed by flow cytometry using an anti-FimH antibody) were then incubated with a titration of vaccinated mouse sera and controls. Sera were diluted in PBS + 0.1 % BSA and titrated 2.5-fold for 7 points, starting at 1:100. After 45 minutes at 37 ⁰C, the mixture was added to the plate and incubated for 45 minutes at 37⁰C before washing away any unbound cells. A titration of anti-FimH_LD_ rabbit sera was used as an internal control on every plate. Specificity of bacterial binding to mannan was established by the inclusion of Methyl α-D-mannopyranoside (Sigma) as a negative control, which reduced binding by >95% at 50 mM levels. Adherent cells were measured with a luminescent probe BacTiter Glo (Promega) and read on a Clariostar Plus plate reader. IC_50_ inhibition values were interpolated using sigmoidal dose response variable-slope curve fitting (Graphpad Prism). Titers are the reciprocal of the serum dilution at which half-maximal inhibition is observed. Responders were defined as those with ≥50% inhibition at the starting dilution, had a defined IC_50_, a positive hillslope, r^2^≥0.80 and at least two points trending towards binding inhibition. The limit of detection (LOD) is designated as a dilution of 50, which is half of the maximum dilution. The statistical significance (*p-*value) of differences in responses between groups was determined using an unpaired t-test with Welch’s correction applied to log-transformed data.

An alternative version of the assay was used for only Figure 1 and involved detection using a directly labelled anti-O25b-AF488 antibody and an O25b clinical *E. coli* isolate PFEEC0547 (collected as part of the ATLAS surveillance program).

### Animal studies

Animal studies were conducted according to Pfizer local and global Institutional Animal Care and Use Committee (IACUC) guidelines at an Association for Assessment and Accreditation of Laboratory Animal Care (AAALAC) International-accredited facility.

### FimH murine immunogenicity studies

6–8-week-old female CD-1 mice were obtained from Charles River Laboratories. For each group of mice, 20 animals were immunized three times subcutaneously with 10 µg FimH protein mixed with 20 µg Quillaja Saponaria-21 (QS-21) from a 5.1 mg / ml QS-21 stock solution containing 5 mM Succinate, 60 mM NaCl, 0.1% PS80, pH 5.6. Mice were bled 2 weeks following immunization. Blood was withdrawn in 3.5-mL serum tubes at each time point and spun in a centrifuge at 3,000 rpm for 10 min. The serum fractions were collected and stored in cryovials.

### Anti-FimH Monoclonal antibody production

CD-1 mice immunized with purified WT FimH-DSG or FimH-DSG V27C L34C proteins in combination with QS21 adjuvant described above received a final intraperitoneal boost of 10 µg mixed with 20 µg QS21 FimH protein 4 days before fusions. Spleens cells from mice with high titers were harvested and fused with the myeloma P3X63-Ag8.653 cell line using polyethylene glycol (P7306, SIGMA HYBRI-Max). Fused cells were cultured in 96 well plates at 37 °C, 8 % CO_2_ in DMEM containing HAT supplement (21060-017 Gibco). After 10 days in culture, hybridomas were screened by enzyme-linked immunosorbent assay (ELISA) using Maxisorp high binding 96 well plates (442404, Thermo Fisher Scientific) coated with 100 ng of FimH-DSG WT protein. Positive hybridomas secreting anti-FimH antibody were subcloned and clonality was assessed by sequencing. Over 300 parent hybridoma Mabs were screened for the ability to inhibit *E. coli* binding to yeast mannan using a single point titration assay. Following clonal expansion and competitive binding (binning) experiments (described in the supplemental material), 5 groups of antibodies with non-overlapping binding sites were identified.

### FimH-DSG TM crystallization and structure determination

Purified FimH-DSG TM, with triple stabilizing mutations (G15A, G16A, V27A), aglycosylation mutations (N7S, N70S, N228Q), a 7-residue Gly-Ser linker, FimG donor strand peptide and a C-terminal 8xHis tag, was buffer exchanged with 20 mM Tris (pH 7.5) using PD-10 column and concentrated to 10 mg/ml. Crystallization was performed at 20 °C using sitting drop vapor diffusion method by mixing equal volumes of protein and reservoir solution containing 1 M Sodium Acetate (pH 4.5) and 25% (w/v) PEG 3350. Crystals grew to their maximum size in ∼7 days. Crystals were cryoprotected using the reservoir solution supplemented with 15% glycerol and flash frozen in liquid nitrogen. Diffraction data were collected at APS 17-ID. The data was processed using autoPROC (Global Phasing Limited) and the structure was solved using WT FimH-DSG complex (PDB code 4XOD) as a starting model. Model building and refinement were carried out using COOT (67) and BUSTER (Global Phasing Limited).

### Characterization of ability of Mab to target ligand binding site

This experiment was conducted on OCTET HTX using Ni-NTA biosensors and buffer containing 1x PBS, 1 % BSA and 0.0 5% Tween 20. Two-fold dilutions (from 10 µg/ml to 0.078125 µg/ml) of Octyl-mannopyranoside ligand were prebound to 5 µg/ml of His-tagged FimH_LD_ WT for 10 min. Biosensors were allowed to capture FimH_LD_ WT prebound to ligand for 5 min. The baseline was established before letting the biosensors loaded with FimH and different concentrations of ligand allowed to bind 5 µg/ml FimH Mab for 5 min. Nanometer response of antibody binding obtained from each dilution of the ligand was plotted against ligand concentration.

### Cryo-EM Sample Preparation, Data Collection, and Processing

For cryo-EM studies, Fab fragments of Mabs 299-3, 304-1, 329-2, 440-2 and 445-3 were generated using one of two methods 1) using a Mouse IgG1 Fab and F(ab′)^2^ Preparation Kit (Thermo Scientific™ Pierce™) which uses immobilized Ficin for cleavage 2) production of recombinant Fab fragments with a C-terminal his tag in ExpiCHO cells, by LakePharma. Cryo-EM sample preparation, data collection and processing are described in full in the supplemental methods.

### Statistical analysis

Statistical significance (*p-*value) of differences in responses between mouse groups was determined using an unpaired t-test with Welch’s correction, applied to log-transformed data.

## Acknowledgements

Thank you to Elliot Dean and Lily Liu for molecular biology and thermal shift support. Thank you to the Pfizer Pearl River Comparative Medicine group for their support for animal studies. We thank and Andy Weiss for critical review of the manuscript. We thank Christina D’Arco for scientific writing support.

## Author Contributions

Conceptualization: YC, MCG, NCS, RGKD, LC Methodology: LC, YC, SS, JL, JH, SK, MCG

Investigation: DK, LC, JL, CP, JA, TC, AI, SK, LL, AG, AE, JH, JM, CK, MK, KC, SS, MCG

Visualization: JJ, JL, HW, YC, NCS

Project administration: NCS, RGKD, ASA

Supervision: NCS, RGKD, YC, AA

Writing – original draft: NCS, LOC, YC

Writing – review & editing: HW, JJ, YVM, MCG, JL, JH, SK, CP, AE, LL, DK, JA, TC, AI, CK, MK, KC, SS, AG, AA, RGKD

## Data and materials availability

All data associated with this study are present in the paper or the Supplementary Materials.

## Supplementary Materials

### 1. Supplementary Methods

#### 1.1. X-ray crystallography of FimH-DSG TM

For X-ray crystallography experiments, FimH-DSG TM was expressed in ExpiCHO cells (Thermo Fisher Scientific) as secreted proteins with C-terminal His tags. Cell culture supernatant was harvested and 1 M Tris pH 7.4 and 5 M NaCl were added to final concentrations of 20 mM and 150 mM final concentrations respectively. A 5 kDa TFF cassette buffer was rinsed and equilibrated in 20 mM Tris pH 7.5 with 500 mM NaCl and 40 mM imidazole. Supernatant was concentrated 2-fold and diafiltered against 6 volumes of 20 mM Tris pH 7.5 500 mM NaCl 40 mM imidazole. Retentate was collected and filtered with a 0.2 um bottletop filter. An XK26/20 column was packed with Ni-Sepharose 6 Fast Flow resin (Cytiva Life Sciences) and equilibrated with 5 column volumes of 20 mM Tris pH 7.5 500 mM NaCl 40 mM imidazole. Retentate was applied at half flow rate and washed until a stable baseline was reached (approximately 55 column volumes). Bound protein was eluted with 20 mM Tris, 500 mM NaCl, 500 mM imidazole, pH 7.5. Fractions containing the protein of interest were pooled and dialyzed in a 2 kDa dialysis cassette against 20 mM sodium acetate, pH 4.3 at 4 °C with two buffer changes. Protein was applied to a SP-Sepharose cation exchange column (Cytiva Life Sciences) that had been equilibrated with the same buffer. Material bound to the cation-exchange resin was eluted with a linear gradient of NaCl using 20 mM sodium acetate, pH 4.3, 1 M NaCl buffer. Fractions were pooled, and dialyzed against TBS, pH 7.4.

Purified FimH-DSG TM was buffer exchanged against 20 mM Tris (pH 7.5) using PD-10 column and concentrated to 10 mg/ml. Crystallization was performed at 20 °C using sitting drop vapor diffusion method by mixing equal volumes of protein and reservoir solution containing 1 M sodium acetate (pH 4.5) and 25% (w/v) PEG3350. Crystals grew to their maximum size in ∼ 7 days. Crystals were cryoprotected using the reservoir solution supplemented with 15% glycerol and flash frozen in liquid nitrogen. Diffraction data were collected at APS 17-ID. The data was processed using autoPROC (Global Phasing Limited) and the structure was solved using WT FimH-DSG complex (PDB code 4XOD) as a starting model (18). Model building and refinement were carried out using COOT and BUSTER (Global Phasing Limited).

#### 1.2. Binning of monoclonal antibodies

Of the 300 parents Mabs screened in the *E. coli* binding inhibition assay, 34 parents were inhibitory; hybridomas from these parents were expanded and 26/34 survived the expansion. From these, 30 ml supernatant was purified using protein A/G resin on gravity flow columns. 12 of the 26 parents had inhibitory activity, evaluated in a 7-point titration assay starting at 75 nM with 2-fold serial dilutions. These hybridomas were cloned, and 3 clones of each were screened again for neutralizing activity. This led to the selection of independent 10 Mab clones with neutralizing activity (**Table 1**).

Epitope binning and kinetics experiments were performed on an Octet HTX instrument. For each epitope binning experiment, Ni-NTA Biosensors (Sartorius Cat# 18-5103) were pre-wet in assay buffer containing 1x PBS 1% BSA 0.1 % tween 20 for at least 10 minutes. To establish the initial baseline, Ni-NTA biosensors were immersed in assay buffer for 60 sec. His-tagged mammalian FimH_LD_ WT (mFimH_LD_ WT) was loaded at 5 ug/ml for 300 sec onto the baseline-established biosensors. mFimH_LD_ WT-loaded and baseline established biosensors were allowed to bind first Mab at 5 ug/ml for 5 min. the baseline was re-established in assay buffer for 3 min. Finally, the first Mab-loaded and baseline-established biosensors were allowed to bind competing Mab (5 µg / ml) for 300 seconds. The nm shift response was measured for all the antibodies under investigation. Antibodies competing against each other should have low binding response whereas the Mabs binding to different epitopes results in higher binding.

To measure kinetic measurements of different FimH Mabs, anti-mouse IgG Fc (AMC Biosensors, cat# 18-5090) were pre-wet in assay buffer for at least 10 min. The initial baseline was established with pre-wet AMC biosensors in assay buffer for 60 seconds. FimH monoclonal antibodies at 1 µg/ml were loaded onto baseline-established biosensors until the nm threshold reached 1 nm, or for 600 seconds. The baseline was re-established for Mab-loaded biosensors in assay buffer, for 3 minutes. Mab-loaded and baseline-established biosensors were allowed to bind 2-fold dilutions of mammalian derived FimH_LD_ WT (starting at 100 nM), for 300 seconds followed by dissociation for 25 min. Results were processed in Data Evaluation v11.1 HT, by subtracting the sensorgram of 0 nM mammalian WT FimH_LD_ blank from the rest of the dilutions. All sensorgrams were aligned to the baseline step followed by aligning the dissociation and association steps. Curves were fit with a Langmuir 1:1 global fit to obtain kinetic measurements. Average values obtained from 2 to 3 experiments were tabulated.

#### 1.3. Sample Preparation for FimH-DSG Fab complex Cryo-EM

To prepare FimH DSG complex with Fabs for analysis by cryo-EM, FimH DSG was combined with a 1.25-fold molar excess each of ECO-329-2 Fab and ECO-445-3 Fab and incubated for 2 hours on ice. The mixture was fractionated by size-exclusion chromatography on a Superdex 200 5/150 GL gel filtration column pre-equilibrated in 20 mM HEPES pH 7.5, 150 mM NaCl. Fractions corresponding to the ternary complex were pooled and concentrated to 0.25 mg/ml in a 3 kDa NMWCO Amicon centrifugal ultrafiltration concentrator. This sample was used for grid vitrification for cryo-EM.

Prior to vitrification, 0.3% β-octylglucoside was added to the sample. The sample was then subjected to centrifugation at 13,200 x *g* for 10 min to remove large aggregates. Gold Quantifoil R1.2/1.3 200 mesh grids were made hydrophilic by glow discharge in residual air at 15 mA for 30 seconds using a Pelco Easiglow. Grids were vitrified using a Vitrobot Mark IV at 4°C and 100% humidity. 4 μl of the sample supernatant was applied to Quantifoil Au 200 mesh R1.2/1.3 grids glow discharged in residual air, then blotted from both sides before plunge-freezing in liquid ethane. Grids were stored under liquid nitrogen until imaging.

#### 1.4. Cryo-EM Data Collection and Processing

Grids were imaged in a Titan Krios G2 transmission electron microscope operated at 300 kV equipped with a Falcon 4i direct electron detector and Selectris X imaging filter. All screening and data collection were performed in EPU (Thermo Fisher Scientific). Movies in EER format were collected at 215,000x magnification (0.59 Å magnified pixel size at the specimen level) with a total electron dose of 50 e^-^/ Å^2^. A dataset of 7,584 movies was collected. Movies were subjected to patch motion correction (nominal pixel size = 0.59 Å, EER fractionation into 40 frames, without upsampling) and patch CTF correction in CryoSPARC 3.3.1. Manual particle picking was used to pick 348 particles, which were subjected to 2D classification to yield templates for template-based particle autopicking. Template-based autopicking against the full dataset yielded 861,802 particles, which were extracted in 500-pixel (29.5 nm) boxes Fourier-cropped to 250 pixels (1.18 Å/pixel) and subjected to multiple rounds of 2D classification to remove damaged particles or non-particle picks from the dataset. 199,906 particles were subjected to ab initio modeling in 4 classes. The most well-defined model, comprising 80,368 particles, was subjected to non-uniform 3D gold-standard refinement and reached a resolution of 3.11 Å, but the model quality was compromised by flexibility in the Fc portions of the Fabs and the FimH pilin domain. To improve the model quality, these portions of the map were subtracted using particle subtraction in CryoSPARC, and the subtracted particles were subjected to 3D gold-standard local refinement against the model, using the pose/shift gaussian prior during alignment and cross-validation-optimal non-uniform regularization. The resulting model was resolved to 3.12 Å with significantly improved model quality.

#### 1.5. Model Building and Refinement

To model each of the components, atomic coordinates from the FimH DSG crystal structure reported here and from PDB entries 4U0R, 7C61, 6H3H, and 7DNH for the Fabs (based on their sequence homology to the heavy and light chains of Fabs 329-2 and 445-3, respectively) were rigid-body fitted into the cryo-EM map density. Sequence modifications were made to each of the models to match those of the molecules used in the experiment, then the model was successively hand-built into the map using Coot v0.9.8.1 in alternation with real-space refinement in Phenix v1.20 to produce the final model. The full cryo-EM data processing workflow and validation metrics can be found in the supplementary materials. Figures based on the structure were produced in PyMol v2.5.4, UCSF Chimera v1.16, and ChimeraX v1.4.

### 2. Supplementary data tables

**Table S1.** Bacterial binding inhibitory titers induced by E. coli and mammalian produced FimH proteins in mice.

| **Group** | **Responder rate (%)** | **# of responders (N=20)** | **Binding inhibition /geometric mean IC<sub>50</sub>** |
| --- | --- | --- | --- |
| Periplasmic FimH<sub>LD</sub> WT | 55 | 11 | 300 |
| Periplasmic FimH<sub>LD</sub> V27C L34C | 40 | 8 | 254 |
| Mammalian FimH<sub>LD</sub> WT | 45 | 9 | 194 |
| Mammalian FimH<sub>LD</sub> V27C L34C | 10 | 2 | 73 |
| Mammalian FimH-DSG WT | 75 | 15 | 529 |
| Mammalian FimH-DSG V27C L34C | 80 | 16 | 579 |
| Periplasmic FimCH | 67 | 12 (N=18) | 354 |

**Table S3.**
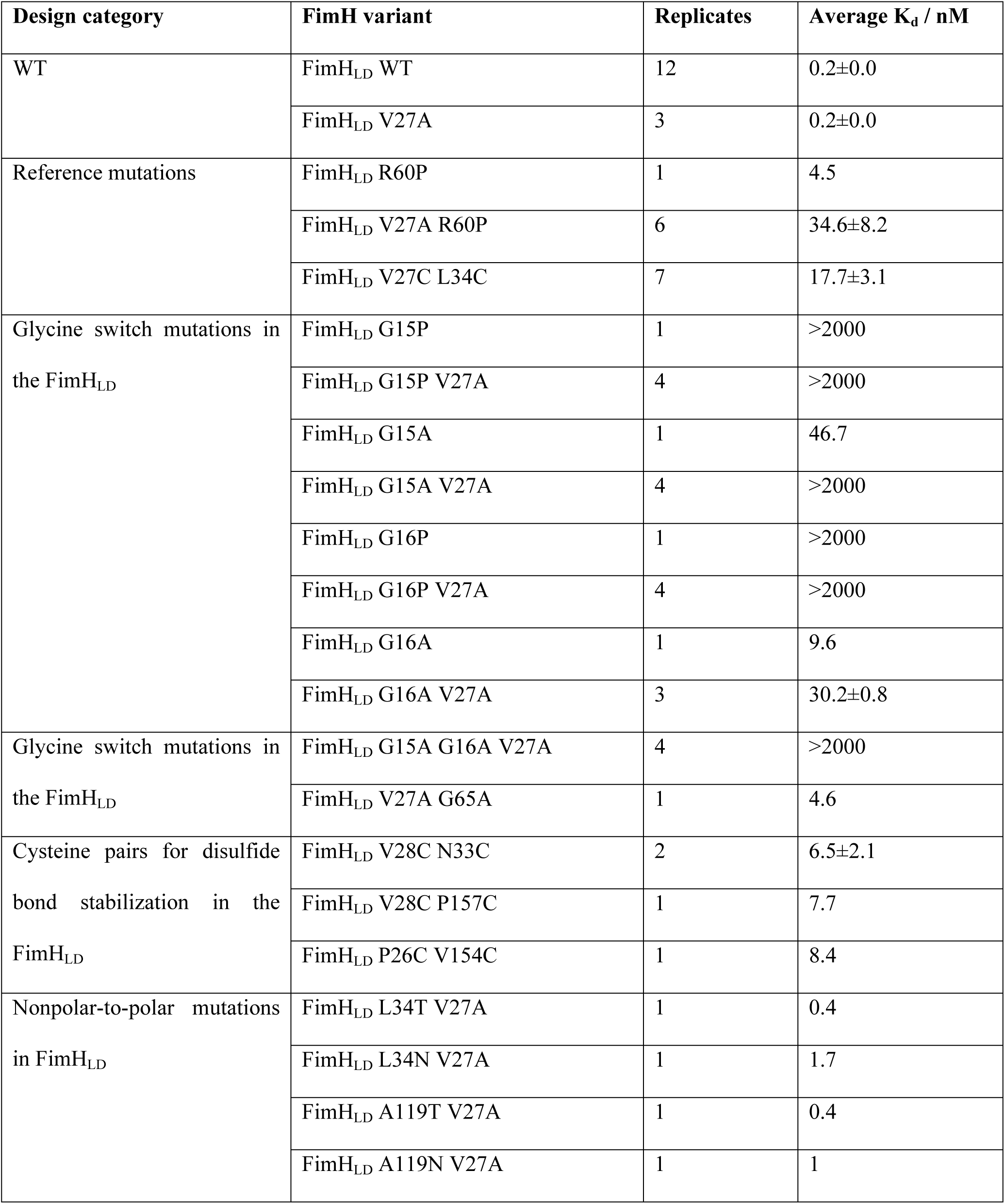
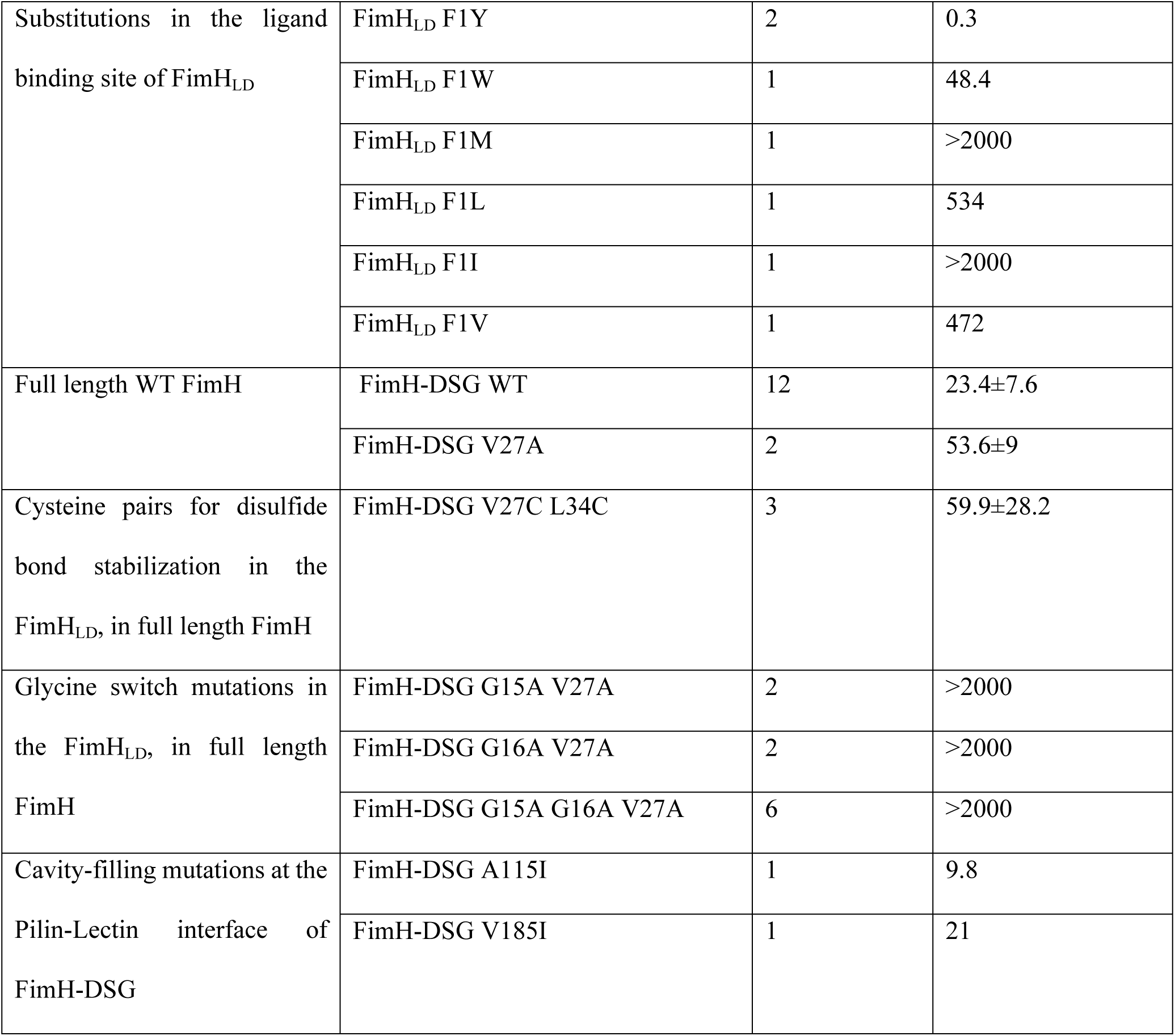
Binding K_d_ of FimH mutants to octylbiphenylmannopyranoside ligand.

**Table S4.**
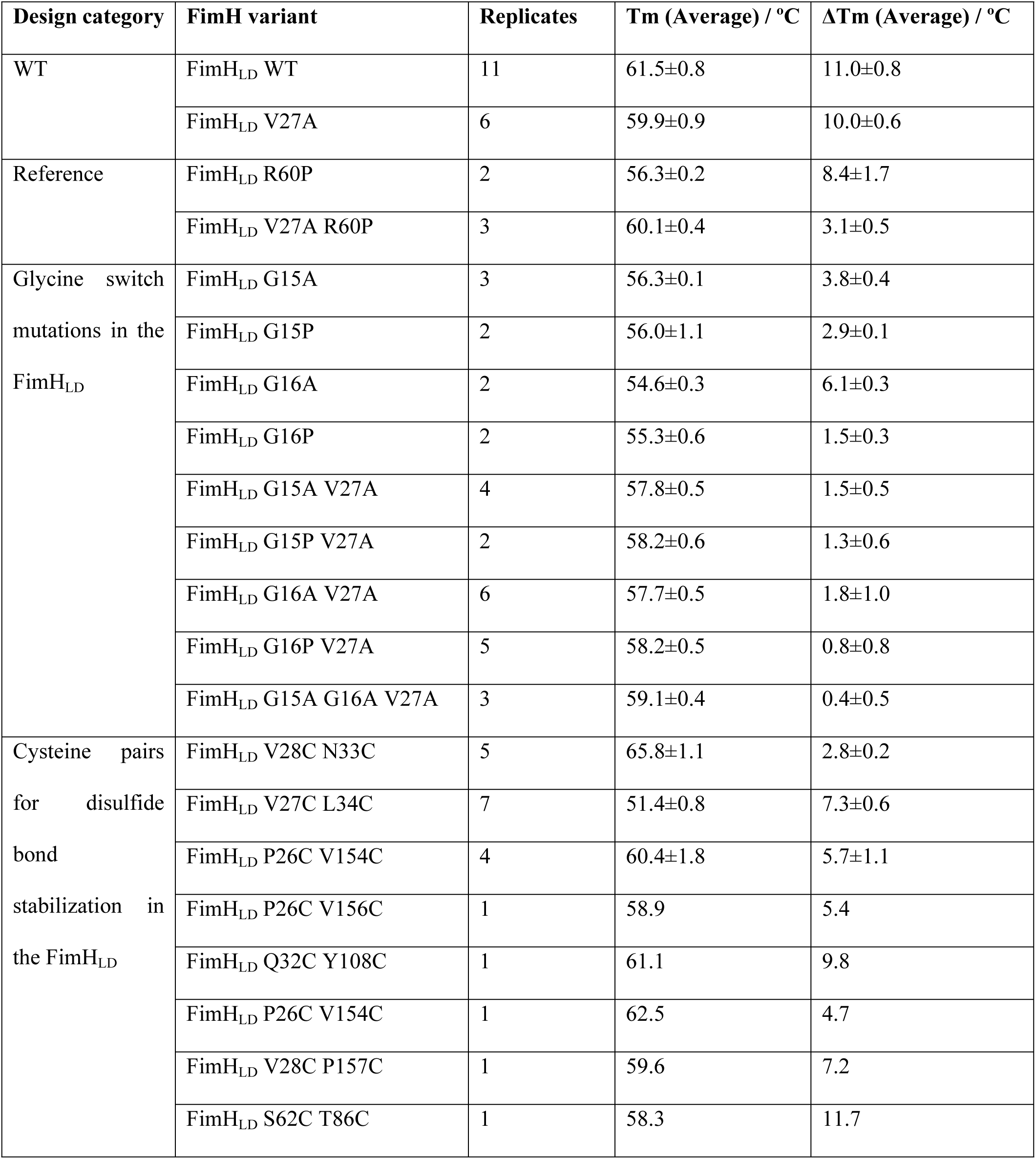
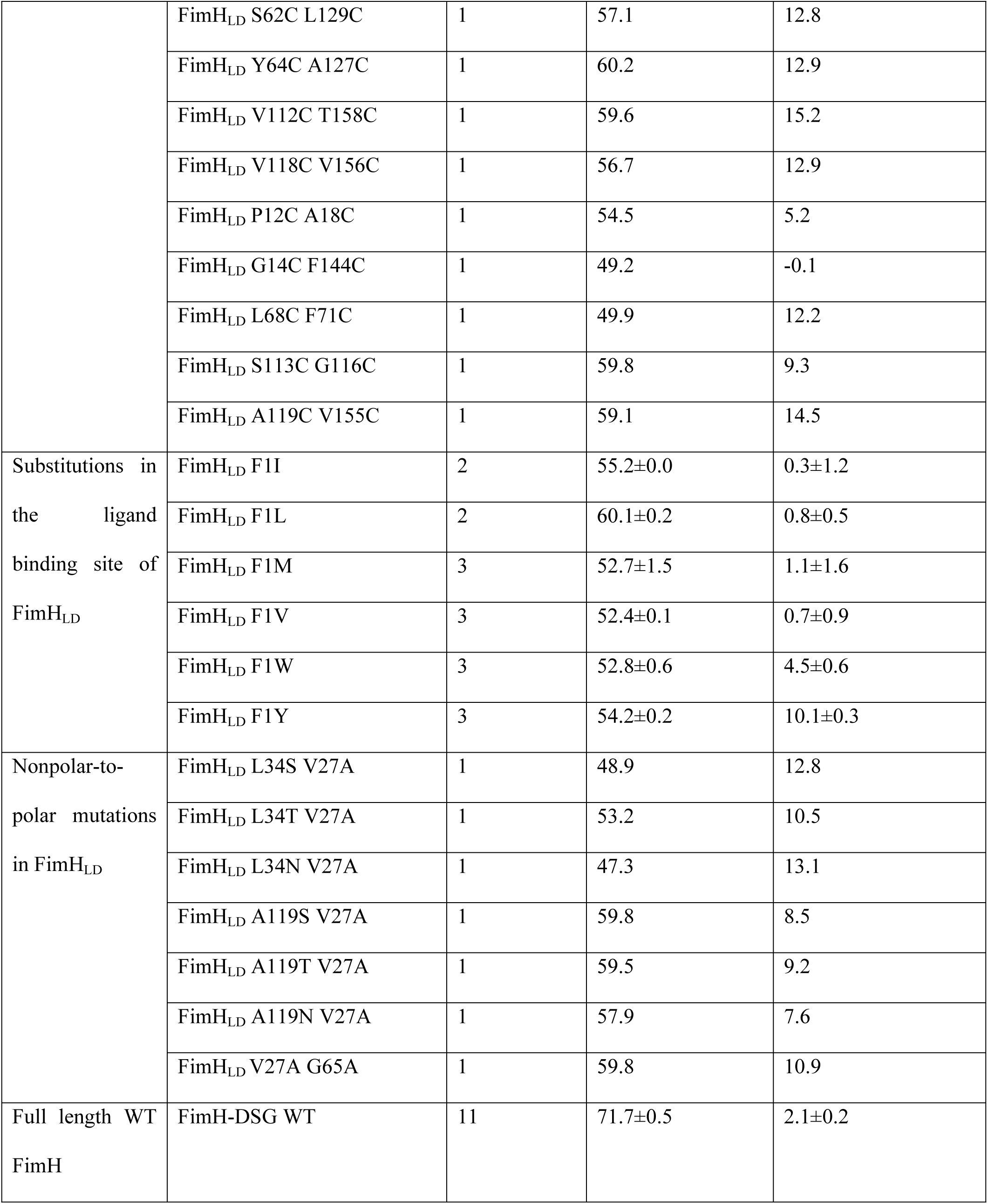
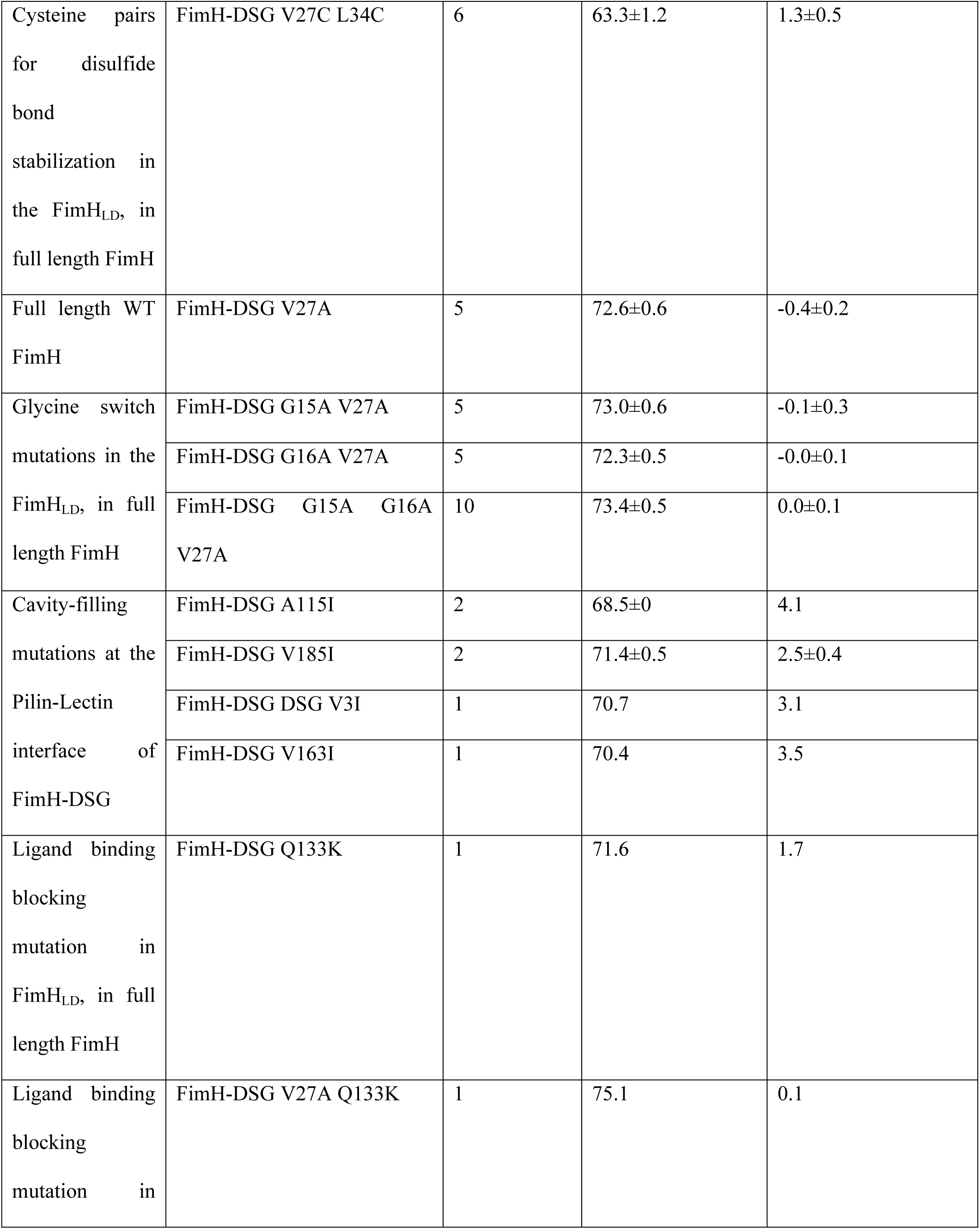
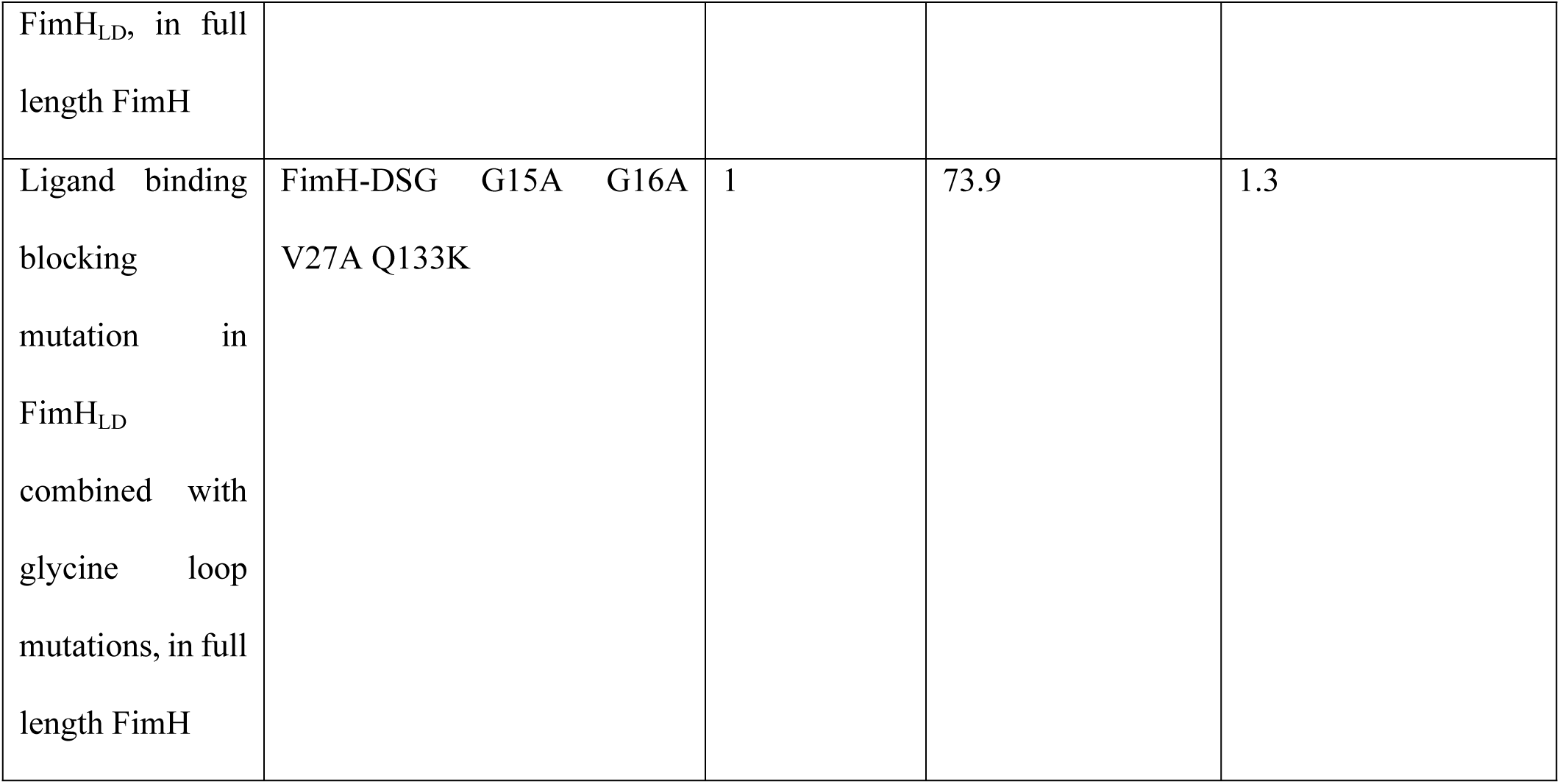
Melting temperature of FimH mutants in apo state and in the presence of methyl alpha-D-mannopyranoside.

**Table S5.** Mab binding to FimH_LD_ mutants.

| FimH variant | Response (nm) |  |  |
| --- | --- | --- | --- |
|  | Monoclonal antibody |  |  |
|  | 299-3 | 304-1 | 440-2 |
| FimH<sub>LD</sub> WT | 3.10 | 3.02 | 0.04 |
| FimH<sub>LD</sub> V27A | 0.44 | 0.54 | 0.06 |
| FimH<sub>LD</sub> V27A R60P | 3.19 | 3.03 | 0.85 |
| FimH<sub>LD</sub> V27A G15A | 3.31 | 3.03 | 0.78 |
| FimH<sub>LD</sub> V27A G15P | 3.07 | 2.95 | 0.62 |
| FimH<sub>LD</sub> V27A G16A | 3.44 | 3.20 | 0.75 |
| FimH<sub>LD</sub> V27A G16P | 3.26 | 3.16 | 0.88 |
| FimH<sub>LD</sub> G15A G16A V27A | 3.34 | 3.06 | 0.79 |
| FimH<sub>LD</sub> V27C L34C | 3.11 | 2.77 | 0.81 |
| FimH<sub>LD</sub> V28C N33C | 3.054 | 2.74 | 0.72 |
| FimH<sub>LD</sub> P26C V154C | 3.11 | 2.89 | 0.83 |

**Table S6.** Mab binding to FimH-DSG variants.

| FimH variant | Response (nm) |  |  |
| --- | --- | --- | --- |
|  | Monoclonal antibody |  |  |
|  | 299-3 | 304-1 | 440-2 |
| FimH-DSG WT | 2.44 | 2.22 | 0.38 |
| FimH-DSG V27A | 2.48 | 2.17 | 0.35 |
| FimH-DSG V27C L34C | 2.35 | 2.23 | 0.33 |
| FimH-DSG V27A G15A | 2.52 | 2.30 | 0.37 |
| FimH-DSG V27A G16A | 2.69 | 2.42 | 0.42 |
| FimH-DSG G15A G16A V27A | 2.68 | 2.30 | 0.49 |

**Table S7.** Bacterial binding inhibitory titers induced by FimH_LD_ and FimH-DSG mutants proteins in mice.

| Protein | IC<sub>50</sub> GMTs |  | Responder rate (%) |  | Responders (n) |  | Mice (n) |  |
| --- | --- | --- | --- | --- | --- | --- | --- | --- |
|  | PD2 | PD3 | PD2 | PD3 | PD2 | PD3 | PD2 | PD3 |
| FimH<sub>LD</sub> WT | 89 | 191 | 20 | 40 | 4 | 8 | 20 | 20 |
| FimH<sub>LD</sub> V27A | 104 | 439 | 26 | 61 | 5 | 11 | 19 | 18 |
| FimH-DSG V27A | 1175 | 6102 | 78 | 100 | 14 | 18 | 18 | 18 |
| FimH<sub>LD</sub> G15A V27A | 57 | 683 | 5 | 53 | 1 | 10 | 20 | 19 |
| FimH-DSG G15A V27A | 1740 | 3400 | 84 | 100 | 16 | 19 | 19 | 19 |
| FimH<sub>LD</sub> G15P V27A | 58 | 346 | 5 | 42 | 1 | 8 | 20 | 19 |
| FimH<sub>LD</sub> G16A V27A | 93 | 1193 | 13 | 69 | 2 | 11 | 16 | 16 |
| FimH<sub>LD</sub> G16P V27A | 91 | 352 | 10 | 45 | 2 | 9 | 20 | 20 |
| FimH<sub>LD</sub> G15A G16A V27A | 111 | 1307 | 26 | 63 | 5 | 12 | 19 | 19 |
| FimH-DSG G15A G16A V27A | 1869 | 2386 | 84 | 95 | 16 | 18 | 19 | 19 |
| FimH<sub>LD</sub> V27A R60P | 212 | 1056 | 32 | 63 | 6 | 12 | 19 | 19 |
| FimH<sub>LD</sub> V28C N33C | 103 | 461 | 16 | 53 | 3 | 10 | 19 | 19 |

**Table S8.**
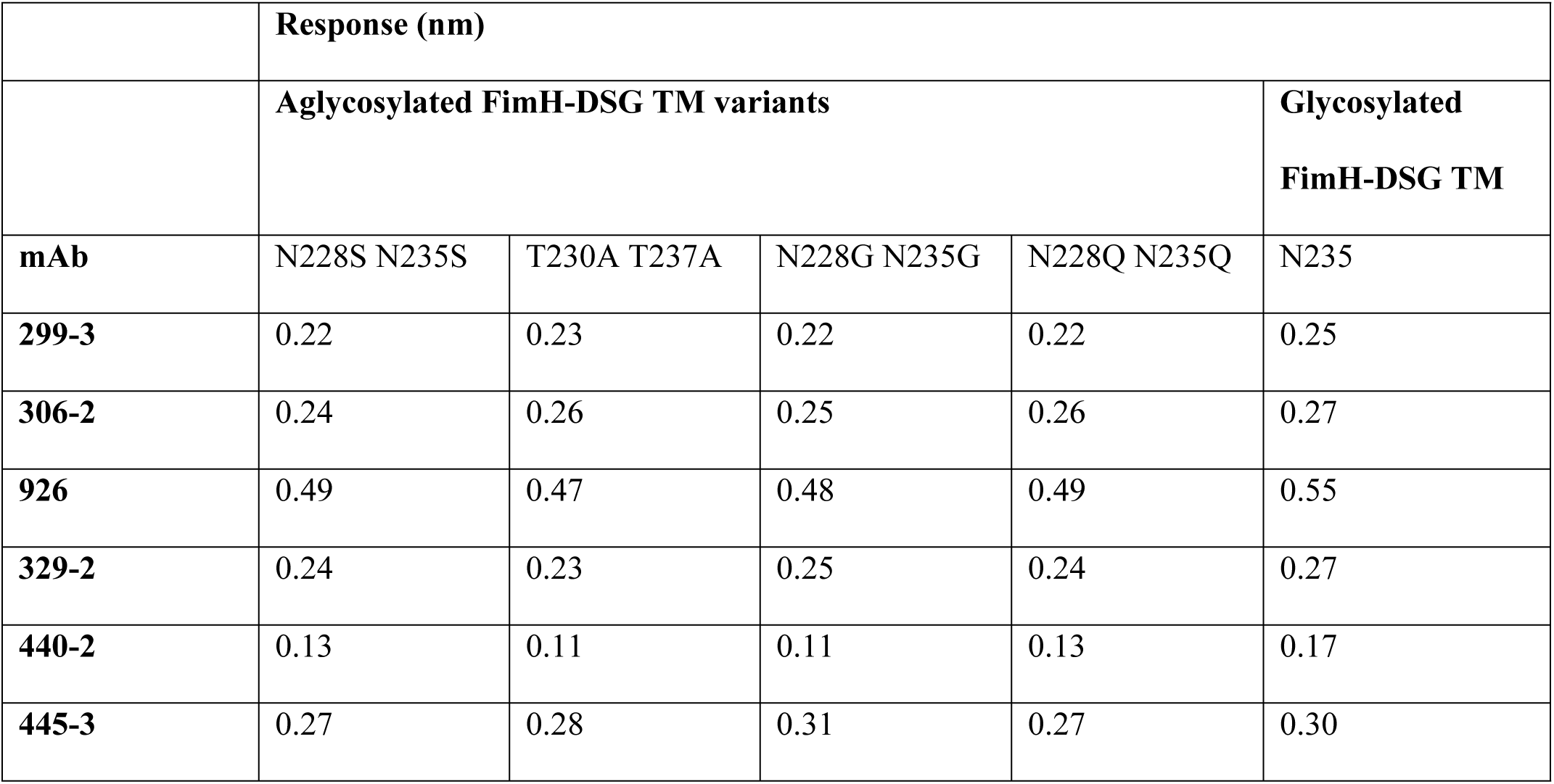
Neutralizing epitopes are preserved in four aglycosylated variants of the FimH DsG triple mutant antigen.

**Table S9.**
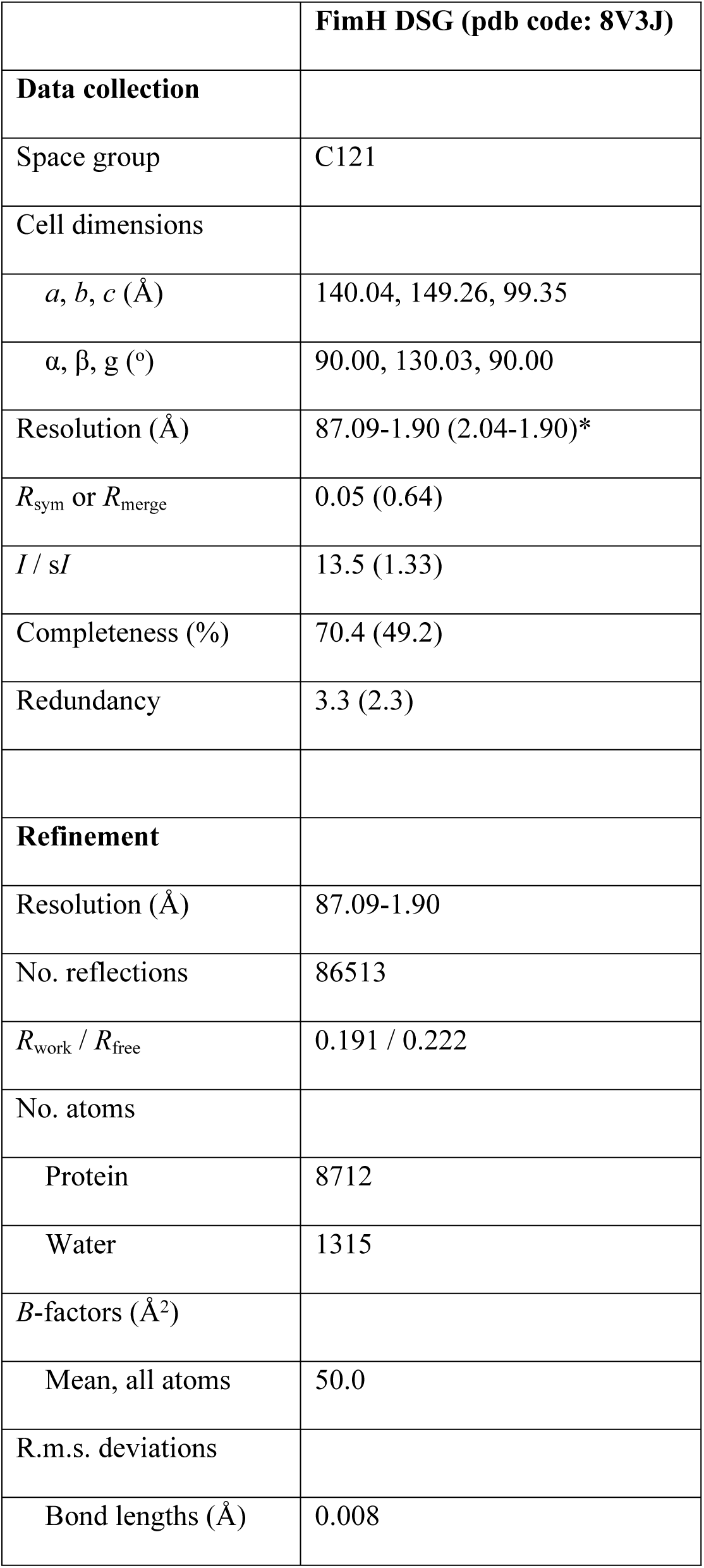
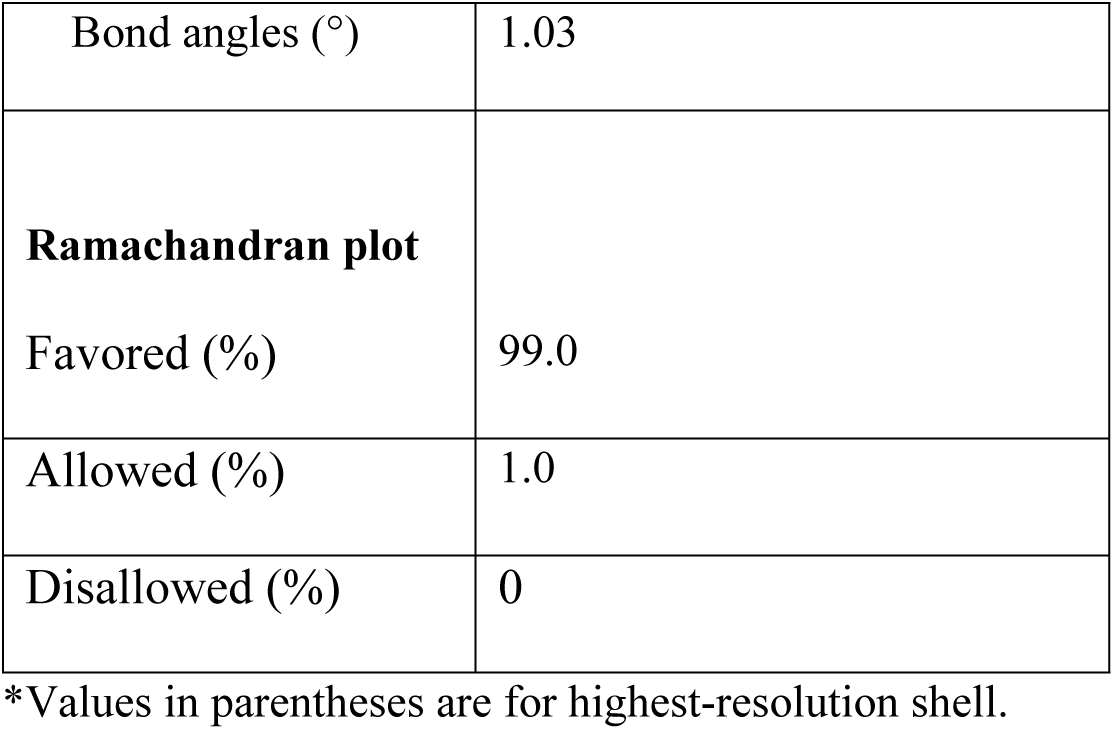
X-ray data collection and refinement statistics for FimH DSG.

|  | **FimH DSG (pdb code: 8V3J)** |
| --- | --- |
| **Data collection** |  |
| Space group | C121 |
| Cell dimensions |  |
| $a, b, c$ (Å) | 140.04, 149.26, 99.35 |
| $\alpha, \beta, \gamma$ (°) | 90.00, 130.03, 90.00 |
| Resolution (Å) | 87.09-1.90 (2.04-1.90)* |
| $R_{\text{sym}}$ or $R_{\text{merge}}$ | 0.05 (0.64) |
| $I / \sigma I$ | 13.5 (1.33) |
| Completeness (%) | 70.4 (49.2) |
| Redundancy | 3.3 (2.3) |
| **Refinement** |  |
| Resolution (Å) | 87.09-1.90 |
| No. reflections | 86513 |
| $R_{\text{work}} / R_{\text{free}}$ | 0.191 / 0.222 |
| No. atoms |  |
| Protein | 8712 |
| Water | 1315 |
| $B$-factors (Å<sup>2</sup>) | |
| Mean, all atoms | 50.0 |
| R.m.s. deviations |  |
| Bond lengths (Å) | 0.008 |
| Bond angles (°) | 1.03 |
| **Ramachandran plot** |  |
| Favored (%) | 99.0 |
| Allowed (%) | 1.0 |
| Disallowed (%) | 0 |

\*Values in parentheses are for highest-resolution shell.

**Table S10.** Cryo-EM data and refinement statistics.

| **Data collection and processing** | **FimH DSG TM + Fab 440-2 + Fab 445-3** | **FimH DSG TM + Fab 329-2 + Fab 445-3**<br>**(EMDB-43048)**<br>**(PDB 8V93)** |
| --- | --- | --- |
| Magnification | 165,000x | 215,000x |
| Voltage (kV) | 300 | 300 |
| Electron exposure (e-/Å<sup>2</sup>) | 50 | 50 |
| Defocus range (μm) | -3.2 to -1.2 | -2.4 to -0.6 |
| Pixel size (Å) | 0.87 | 0.59 |
| Symmetry imposed | C1 | C1 |
| Initial particle images (no.) | 2,817,610 | 861,802 |
| Final particle images (no.) | 199,492 | 80,368 |
| Map resolution (Å) | 3.9 | 3.12 |
| FSC threshold | (0.143) | (0.143) |
| **Refinement** |  |  |
| Initial model used (PDB code) |  | 4U0R, 7C61, 6H3H, 7DNH |
| Model resolution (Å) |  | 3.9 |
| FSC threshold |  | 0.5 |
| Map sharpening *B* factor (Å<sup>2</sup>) |  | 82.7 |
| Model composition |  |  |
| Non-hydrogen atoms |  | 4,677 |
| Protein residues |  | 613 |
| Ligands |  | 0 |
| *B* factors (Å<sup>2</sup>) |  |  |
| Protein |  | 29.58/100.90/56.49 |
| Ligand |  | N/A |
| R.m.s. deviations |  |  |
| Bond lengths (Å) |  | 0.005 |
| Bond angles (°) |  | 0.969 |
| Validation |  |  |
| MolProbity score |  | 1.826.54 |
| Clashscore |  | 0 |
| Poor rotamers (%) |  |  |
| Ramachandran plot |  |  |
| Favored (%) |  | 92.82 |
| Allowed (%) |  | 7.01 |
| Disallowed (%) |  | 0.17 |

### 3. Supplementary figures

**Fig. S1.**
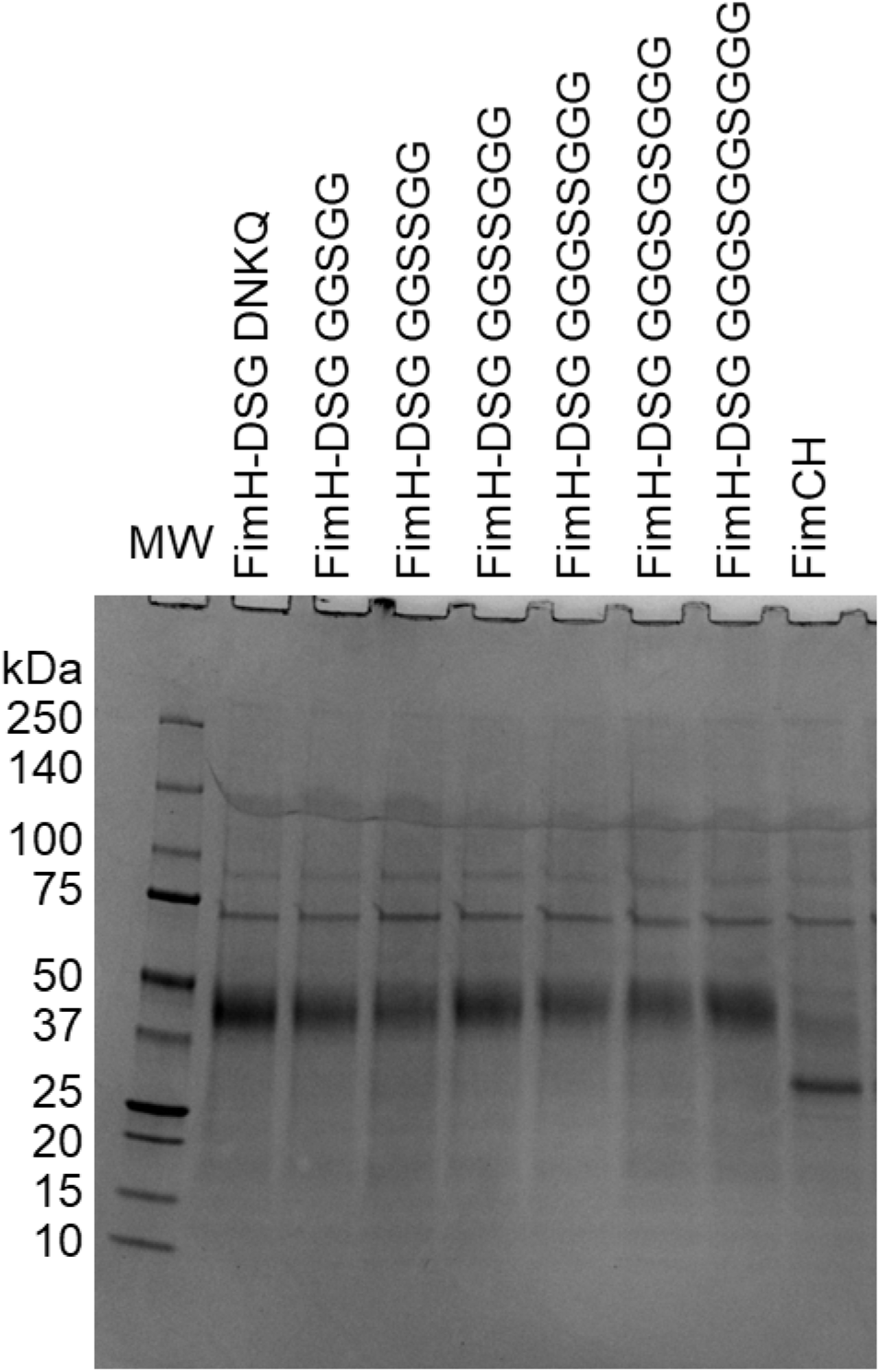
Optimization of full length FimH production in Expi293 cells. Expi293 cells were transfected with constructs encoding FimH-DSG with linkers (DNKQ, GGSSGG, etc.) and the FimCH complex. Raw culture supernatants collected 96 hours post transfection were loaded on an SDS-PAGE gel and stained with Coomassie blue.

**Fig. S2.**
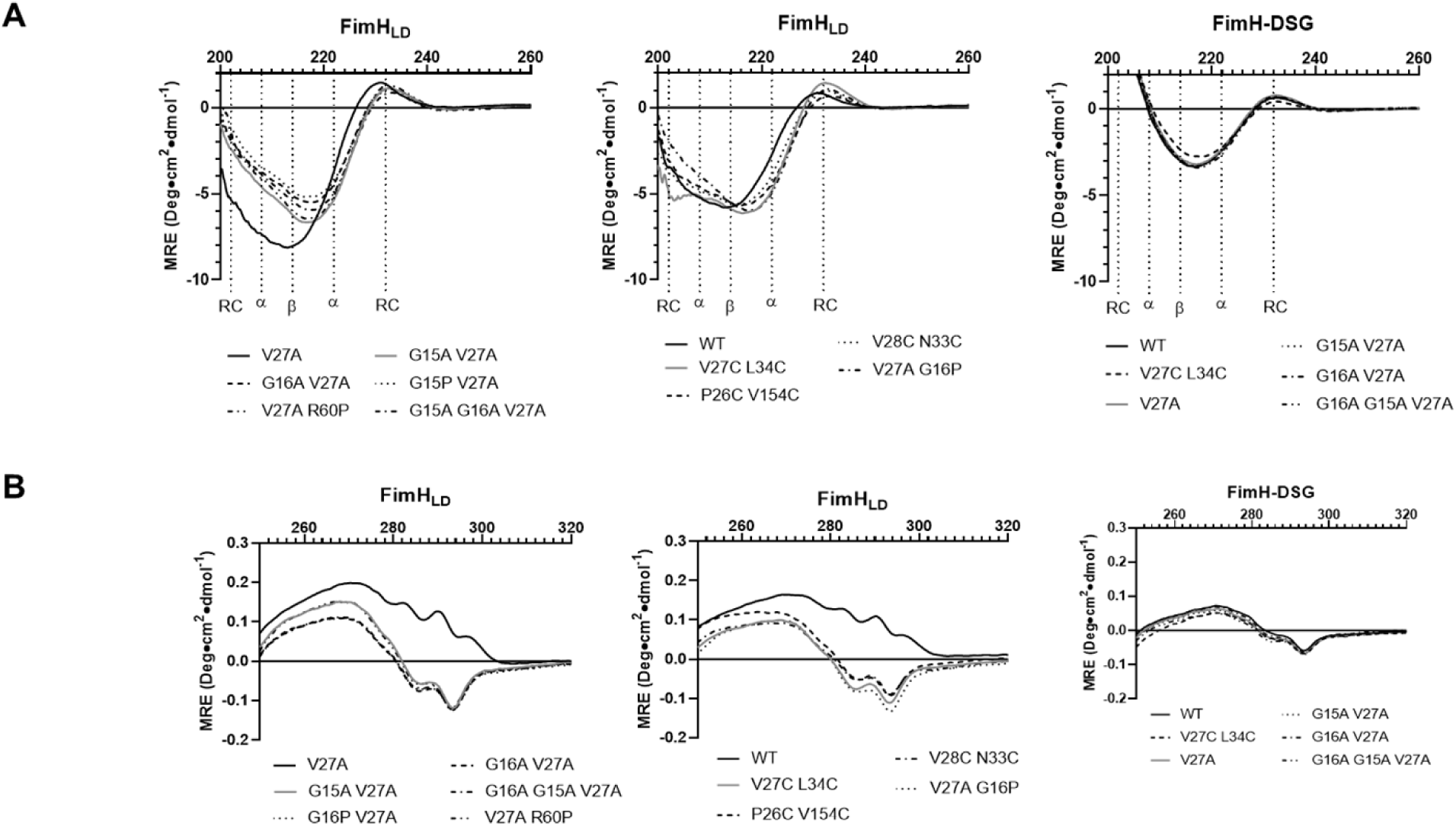
Informative regions of CD spectra of FimH_LD_ and FimH-DSG mutants. CD spectra in the far-UV **(A)**, reporting on the secondary structure, and near-UV **(B)**, reporting on the tertiary structure of FimH variants.

**Fig. S3.**
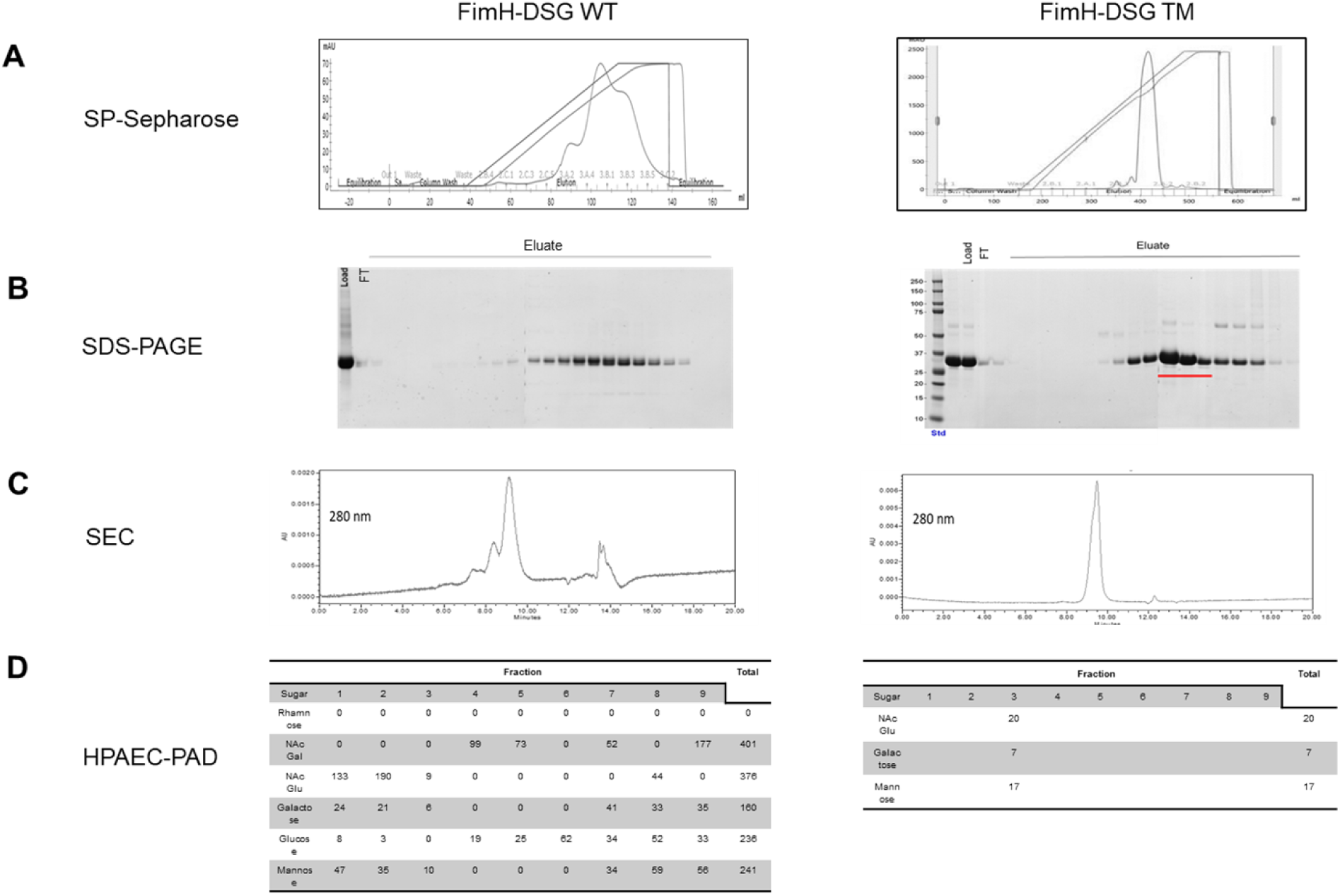
FimH-DSG TM can be purified to homogeneity from ExpiCHO supernatants. To produce protein for structural characterization, FimH-DSG WT and FimH-DSG TM proteins were expressed in ExpiCHO cells as secreted proteins containing C-terminal His tags. (A) Elution profile of FimH-DSG WT (left panel) and FimH-DSG TM (right panel) on SP-Sepharose column. (B) SDS-PAGE analysis of eluted fractions. (C) Analytical SEC of FimH-DSG WT (left panel); wild type FimH-DSG TM (right panel). (D) Normalized amounts of monosaccharides (µg / mg protein) detected by HPAEC-PAD in various SP-Sepharose fractions from FimH-DSG WT and the main peak of FimH-DSG TM.

**Fig S4.**
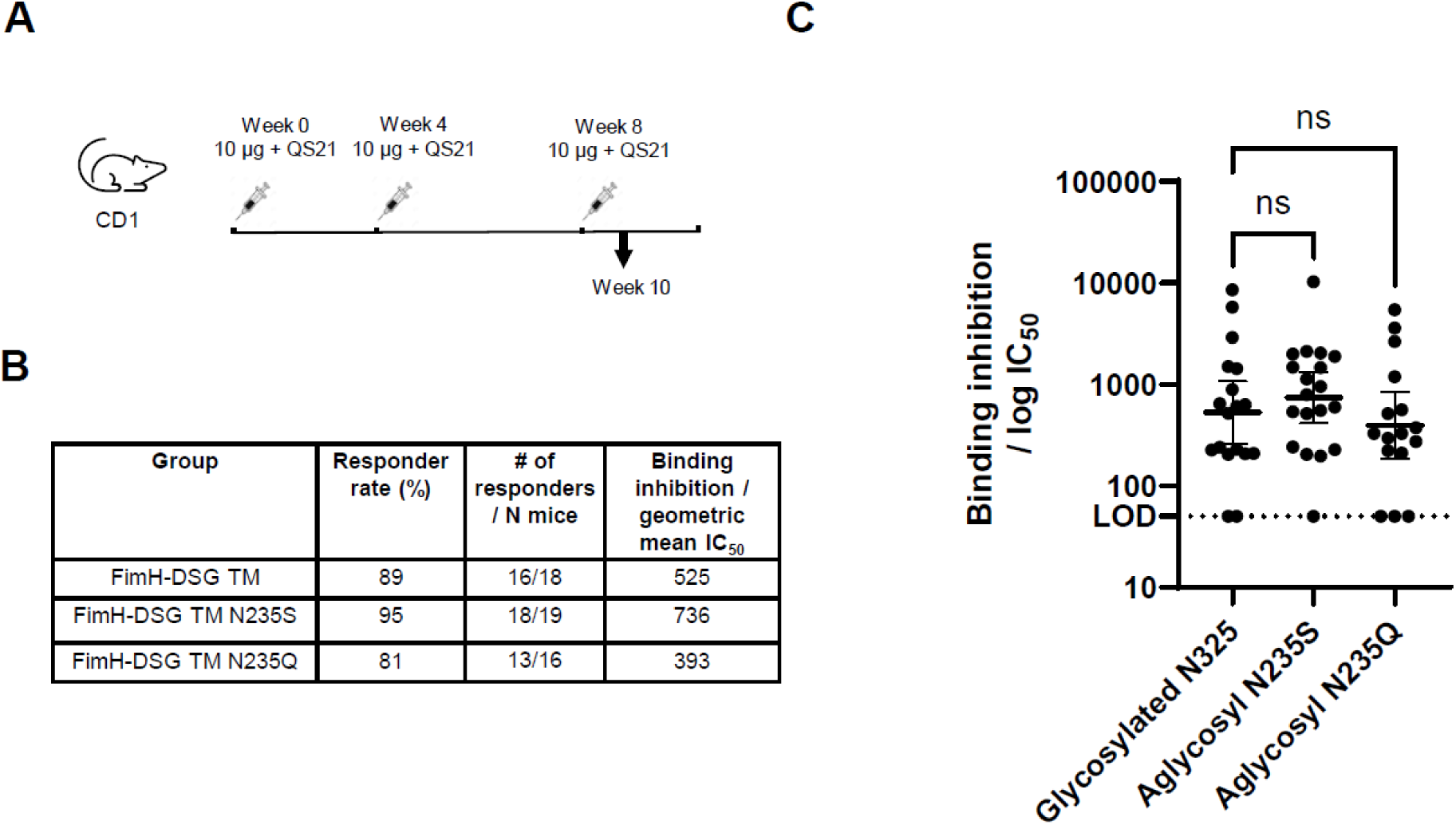
FimH-DSG aglycosylation variants have similar ability to elicit antibodies that inhibit bacterial binding compared to glycosylated parent. **(A)** Groups of 20 CD-1 mice (7-9 weeks old) were vaccinated subcutaneously with 0.1 mL of 10 mg of FimH-DSG antigens adjuvanted with 20 mg QS21/PS80 at weeks 0, 4 and 8. **(B)** Sera from week 6 and week 10 timepoints were tested for activity in the E. coli binding inhibition assay with geometric mean IC_50_ titers reported. Inhibitory titers were determined from serial dilution of sera from vaccinated mice and represent the reciprocal of the dilution of serum at which 50% of bacteria remain bound to the plate and are shown for post dose 3 timepoint. Statistical significance (p-value) of differences in responses between groups was determined using an unpaired t-test with Welch’s correction applied to log-transformed data. Proportion of animals in each group responding to vaccine by exhibiting measurable IC_50_ titers are reported as % responder rates. **(C)** Figure displaying individual IC_50_ values; bars represent geometric mean and 95% confidence interval.

**Fig. S5.**
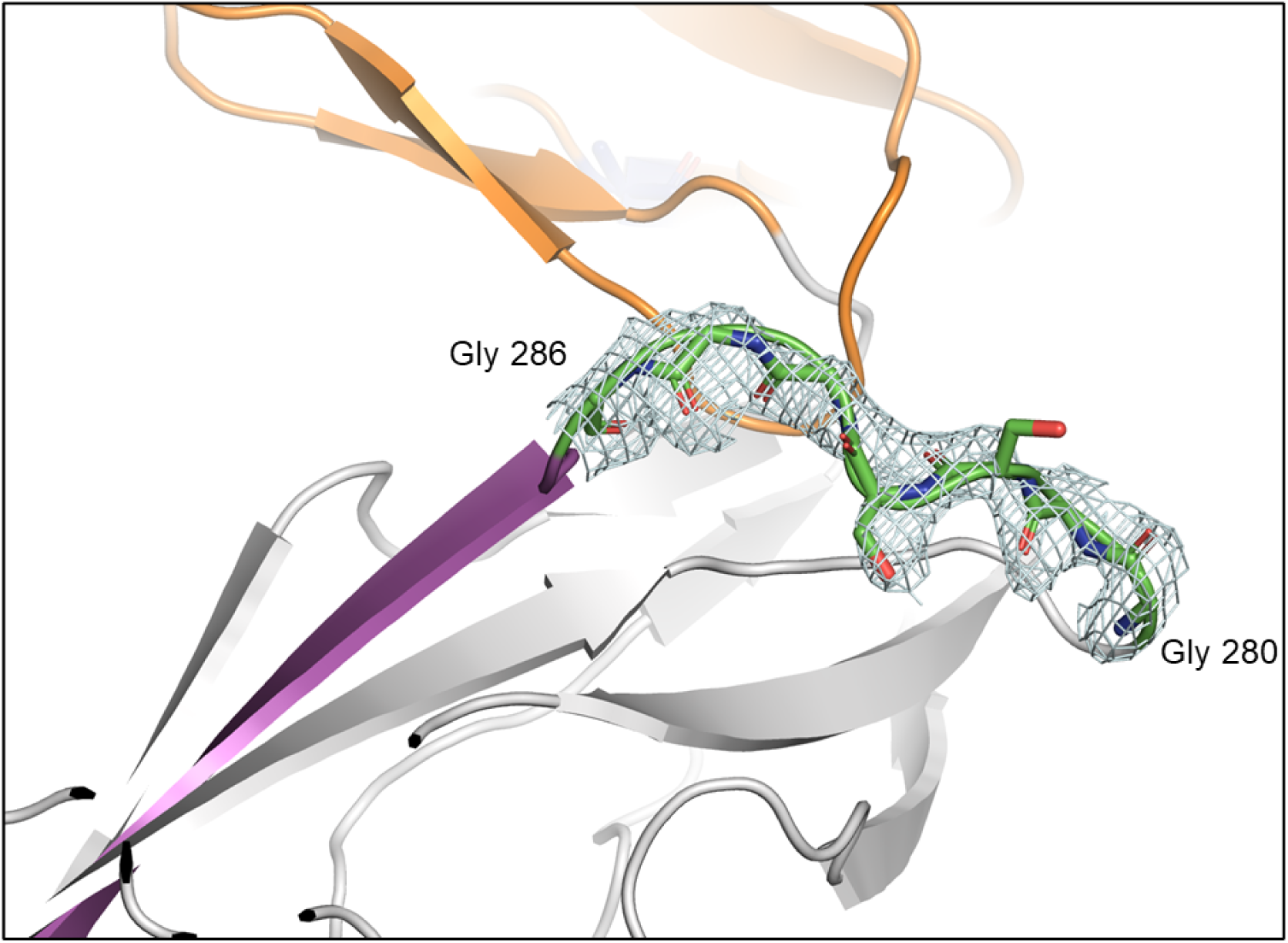
Visualization of linker region in structure of FimH-DSG TM. Electron density of the glycine-serine linker region. FimH_LD_ is shown in orange, FimH_PD_ is shown in grey, the donor-strand peptide in pink and the glycine-serine linker in green.

**Fig. S6.**
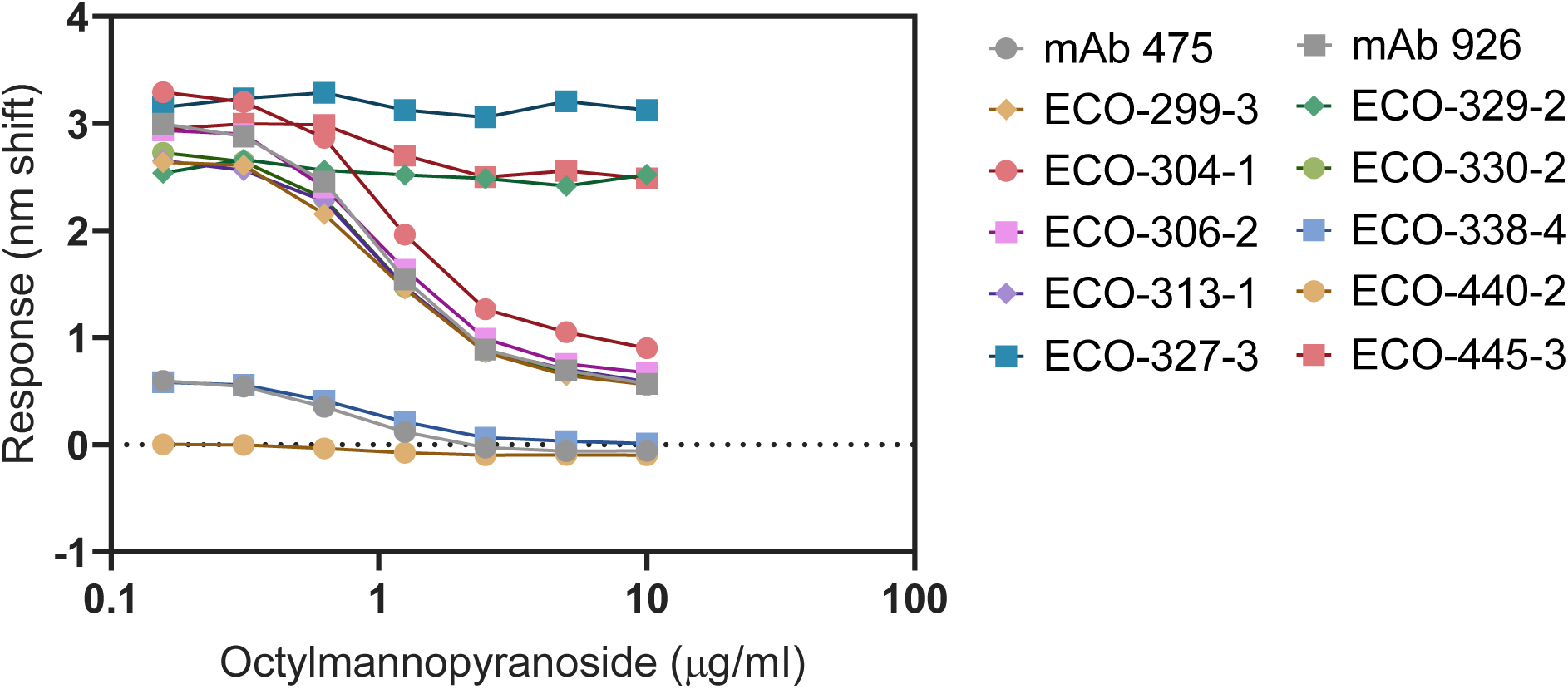
Ability of Octylmannopyranoside to interfere with binding of monoclonal antibodies from FimH_LD_ WT. FimH_LD_ WT was incubated with a two-fold dilution of Ocytlmannopyranoside ligand before loading onto Octet Ni-NTA biosensors. FimH Mabs were allowed to bind at 5 µg/ml to the pre-bound FimH_LD_ and nanometer response measured for each dilution of the ligand.

**Fig. S7.**
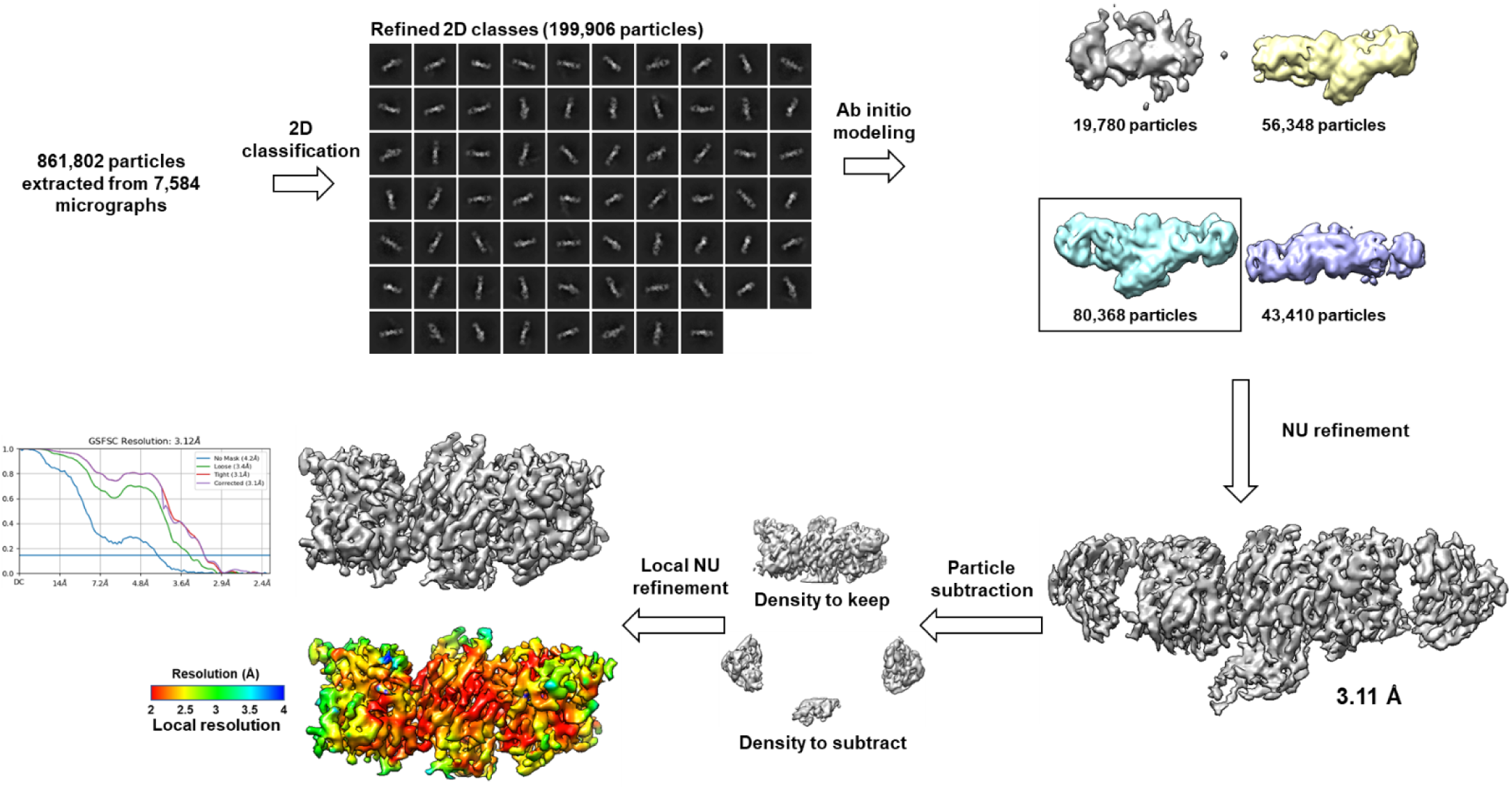
Cryo-EM processing pipeline for FimH/329-2/445-3. Cryo-EM data processing workflow in CryoSPARC for the FimH/329-2/445-3 ternary complex, with final map colored by local resolution and Fourier shell correlation curves from gold-standard refinement.

**Fig. S8.**
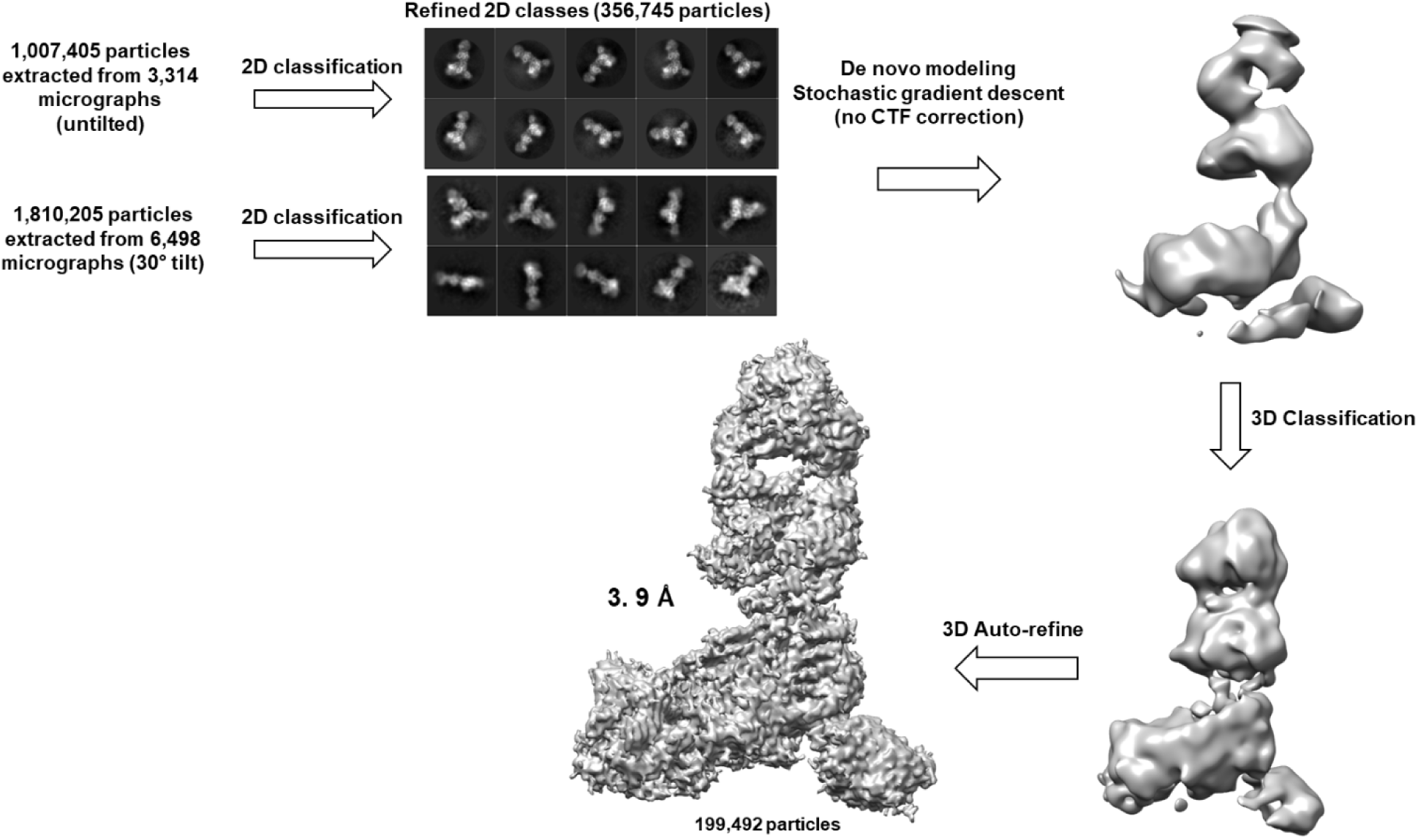
Cryo-EM processing pipeline – FimH/440-2/445-3. Cryo-EM data processing workflow in Relion for the FimH/440-2/445-3 ternary complex.

